# Expansion of a core regulon by transposable elements promotes Arabidopsis chemical diversity and pathogen defense

**DOI:** 10.1101/368340

**Authors:** Brenden Barco, Yoseph Kim, Nicole K. Clay

**Affiliations:** Department of Molecular, Cellular & Developmental Biology, Yale University, Kline Biology Tower 734, 219 Prospect St., New Haven, CT 06511; Hopkins School, 986 Forest Rd, New Haven, CT 06515; Seeds Research, Syngenta Crop Protection, 9 Davis Dr., Durham, NC 27703

## Abstract

Plants synthesize hundreds of thousands of ecologically specialized, lineage-specific metabolites through biosynthetic gene duplication and functional specialization. However, the rewiring of duplicated genes into existing regulatory networks remains unclear. We show that the duplicated gene *CYP82C2* was recruited into the WRKY33 regulon and indole-3-carbonylnitrile (ICN) biosynthetic pathway through exaptation of a retroduplicated LINE retrotransposon (*EPCOT3*) into a novel enhancer. The stepwise development of a chromatin-accessible WRKY33-binding site on *EPCOT3* potentiated the regulatory neofunctionalization of *CYP82C2* and the evolution of inducible defense metabolite 4-hydroxy-ICN in *Arabidopsis thaliana.* Transposable elements (TEs) have long been recognized to have the potential to rewire regulatory networks; these results establish a more complete understanding of how duplicated genes and TEs contribute in concert to chemical diversity and pathogen defense.

---

Plant secondary or specialized metabolites are essential for plant survival in co-evolving biotic and fluctuating abiotic environments. The evolutionary process of chemical innovation resulted in the collective synthesis of hundreds of thousands of ecologically specialized, mostly lineage-specific metabolites (Chae et al., 2014; Weng et al., 2012; Dixon and Strack, 2003; Wink, 2003). Plant specialized metabolic enzymes are ultimately produced from primary metabolic enzymes through gene duplication and subsequent functional divergence of one or both paralogs to produce enzymes with altered expression patterns and/or protein functions (Ohno, 1970; Force *et al*., 1999; Weng *et al*., 2012). They are also often organized into transcription factor (TF) regulons of co-regulated genes for optimal timing, amplitude, and tissue-specific pathway gene expression and subsequent metabolite accumulation (Grotewold, 2005; Hartmann, 2007; Martin *et al*., 2010; Tohge & Fernie, 2012; Omranian et al., 2015).

Changes in *cis*-regulatory modules such as enhancers and promoters can accelerate the capture of duplicated genes into regulons, thus driving phenotypic diversity (Levine and Davidson, 2005; Prud’homme *et al*., 2007; Wray, 2007; Wittkopp & Kalay, 2012; Rogers *et al*., 2013). Enhancers consist of transcription factor binding sites (TFBSs) and are derived either through mutation or co-option of a TFBS-carrying transposable element (TE) (Spitz & Furlong, 2012; Wittkopp & Kalay, 2012). TE exaptations have been hypothesized to be responsible for the rapid transcriptional rewiring of gene regulatory networks in ancient lineages of vertebrates (Feschotte 2008; Bourque 2009; Lynch *et al*., 2011; de Souza *et al*., 2013; Chuong et al., 2016) and plants (Hénaff *et al*., 2014), but general understandings of the physiological significance of this rewiring are greatly limited.

Bacteria elicit two primary immune defense modes in plants, pattern- and effector-triggered immunity (PTI and ETI) (Jones & Dangl, 2006). Pathogenic bacteria additionally compromise PTI via specific virulence effector proteins (effector-triggered susceptibility, ETS; Jones & Dangl, 2006). PTI involves the extracellular perception of conserved molecules known as microbe-associated molecular patterns (MAMPs), whereas ETI involves the cytosolic perception of effectors. Although ETI results in the formation of more rapid and robust pathogen-specific response including the hypersensitive response (HR), a form of programmed cell death (Jones & Dangl, 2006), both result in the ability of naïve host cells to generate, through non-self perception and subsequent transcriptional reprogramming, pathogen-inducible specialized metabolites necessary for defense (Hammerschmidt, 1999; Mansfield, 2000; Clay *et al*., 2009).

Three pathogen-inducible tryptophan (Trp)-derived defense metabolites – camalexin, 4-methoxyindol-3-ylmethylgucosinolate (4M-I3M), and 4-hydroxyindole-3-carbonylnitrile (4OH-ICN) – have been shown to expand innate immunity in *Arabidopsis thaliana* (Bednarek *et al.,* 2009; Clay *et al.,* 2009; Thomma *et al.,* 1999; Tsuji *et al.,* 1992; Rajniak *et al.,* 2015). The three biosynthetic pathways share an early step, which is the conversion of Trp to indole-3-acetaldoxime (IAOx) via the genetically redundant P450 monooxygenases CYP79B2 and CYP79B3 (Fig. 1a) (Hull *et al*., 2000; Mikkelsen *et al*., 2000; Bak *et al*., 2001; Glawischnig *et al*., 2004; Klein *et al*., 2013; Rajniak *et al*., 2015). The camalexin and 4OH-ICN pathways additionally share the conversion of IAOx to indole-3-cyanohydrin (ICY) by partially redundant P450s CYP71A12 and CYP71A13 (Fig. 1a) (Nafisi *et al*., 2007; Klein *et al*., 2013; Rajniak *et al*., 2015). CYP71A13 and CYP71B15/PAD3 catalyze further reactions, leading to camalexin production, whereas the flavin-dependent oxidase FOX1/AtBBE3 and P450 CYP82C2 convert ICY to 4OH-ICN (Fig. 1a) (Nafisi *et al*., 2007; Böttcher *et al*., 2009; Rajniak *et al*., 2015). 4M-I3M is widely distributed across the mustard family (Brassicaceae), whereas camalexin is restricted to the Camelineae tribe of Brassicaceae (Bednarek *et al*., 2011). The evolutionary conservation of 4OH-ICN has not yet been investigated.

**Figure 1.**
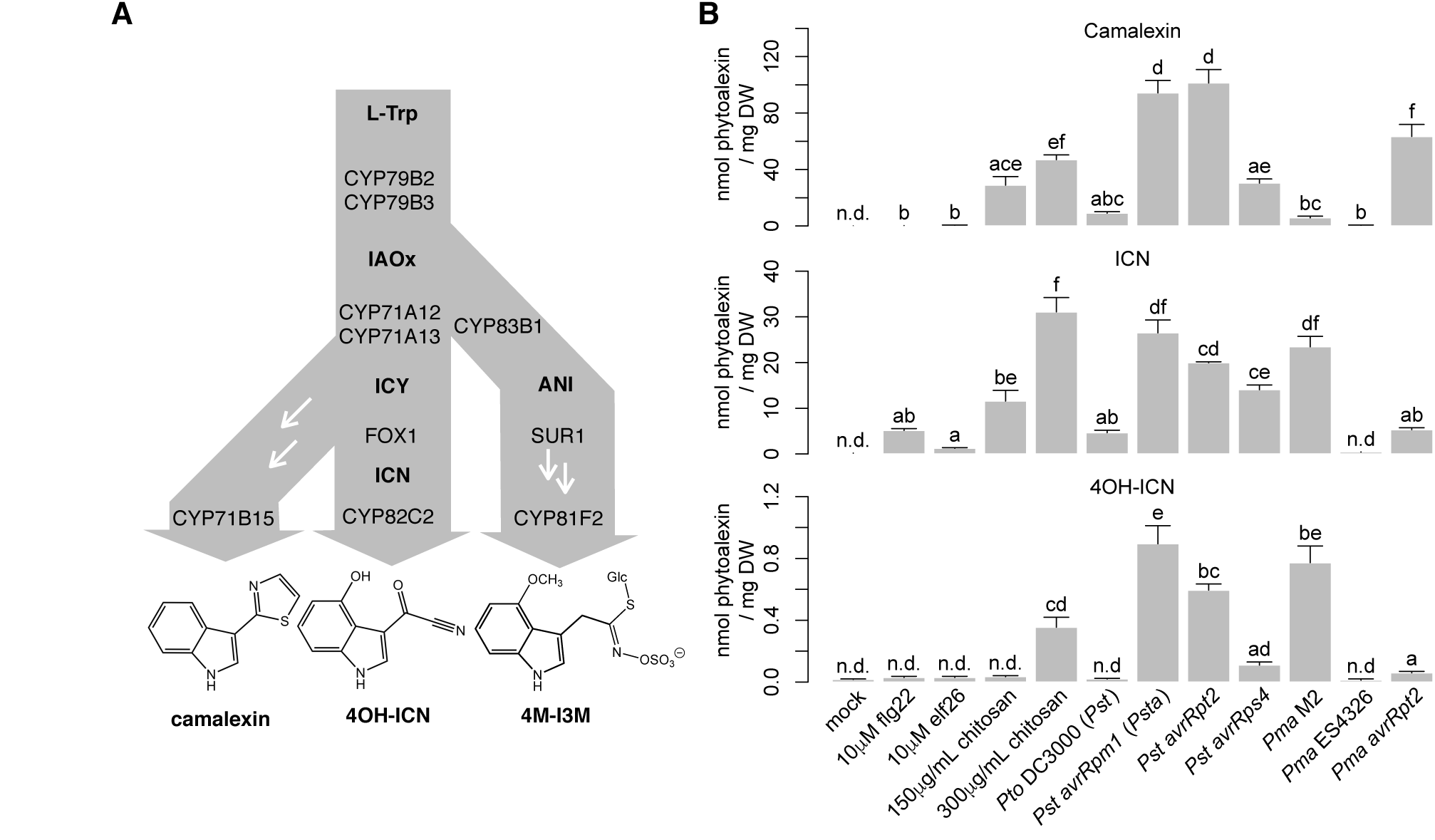
4OH-ICN is synthesized under ETI-like responses. (**a**). Schematic of tryptophan (L-Trp)-derived specialized metabolism in *A. thaliana*. White arrows denote the presence of additional enzymes. ICY, indole cyanohydrin; ANI, *aci*-nitro indole. (**b**). LC-DAD-FLD-MS analysis of camalexin (top), ICN (middle), and 4OH-ICN (bottom) in seedlings elicited with indicated MAMPs and bacterial strains for 27 hr. Data represent mean ± SE of 3-4 biological replicates. Different letters denote statistically significant differences (*P* < 0.05, one-factor ANOVA coupled to Tukey’s test). ICA-ME and 4OH-ICA-ME are methanolic degradation products of ICN and 4OH-ICN, respectively. 4OH-ICA is an aqueous degradation product of 4OH-ICN. For (**b**), source data are provided as a Source Data file.

The TF WRKY33 has been shown to regulate the pathogen-inducible biosynthesis of camalexin in *A. thaliana* and its orthologs regulate numerous unrelated specialized metabolites in other flowering plant lineages (Qiu *et al.,* 2005; Liu *et al.,* 2015; Birkenbihl *et al.,* 2017; Schluttenhofer & Yuan, 2015). The group I class of WRKYs to which WRKY33 belongs is an ancient clade of regulators; orthologs in the green alga *Chlamydomonas reinhardtii* may be ancestral to all higher plant WRKYs (Rinerson *et al*., 2015; Schluttenhofer & Yuan, 2015). While all WRKY TFs bind to the W-box core sequence [TTGAC(T/C)], WRKY33 preferentially binds W-boxes that are within 500 nt of the ‘WRKY33-specific’ motif [(T/G)TTGAAT]) (Rushton *et al.,* 2010; Liu *et al*., 2015).

Here, we show that a recent, lineage-specific TE exaptation resulted in the expansion of a core regulon within the framework of Arabidopsis Trp-derived defense metabolism. Specifically, the LINE retrotransposon *EPCOT3* retroduplicated from a WRKY33-TFBS-carrying progenitor and inserted upstream of the newly duplicated gene *CYP82C2*. Subsequent chromatin remodeling in *A. thaliana* led *EPCOT3* to become a *bona fide* enhancer with demonstrated biochemical, regulatory, physiological, and fitness-promoting characteristics by way of WRKY33-binding and pathogen-responsive *CYP82C2* transcription, 4OH-ICN biosynthesis, and antibacterial defense.

## Results

### 4OH-ICN requires ETI-like responses

To identify the major Trp-derived specialized metabolites synthesized in ETI in *A. thaliana*, we compared host transcriptional and metabolic responses to the PTI-eliciting bacterial MAMPs flg22, elf26, and fungal MAMP chitosan; the PTI/ETS-eliciting pathogens *Pseudomonas syringae* pv. *tomato* DC3000 (*Pto* DC3000 or *Pst*); *Pseudomonas syringae* pv. *maculicola* ES4326 (*Pma*); and the ETI-eliciting pathogens *Pst avrRpm1* (*Psta*), *Pst avrRpt2*, *Pst avrRps4*, *Pma* M2, and *Pma avrRpt2* under similar conditions as those of previous studies (Denoux *et al*., 2008; Clay *et al*., 2009). *Psm* M2 is an ETI-eliciting strain from which the *avrRpm1* gene was originally isolated (Debener *et al*., 1991). Both flg22 and *Psta* induced genes involved in 4OH-ICN, camalexin and 4M-I3M biosynthesis, with 4OH-ICN and camalexin biosynthetic genes having a higher level of induction than those of 4M-I3M in *Psta*-inoculated plants (Supplementary Fig. 1a; Denoux *et al*., 2008). In contrast to the quantitative differences observed in transcriptional responses between PTI and ETI (Tao *et al*., 2003; Navarro *et al*., 2004), the metabolite responses between PTI and ETI differed largely qualitatively. 4M-I3M and the 4M-I3M precursor molecule 4OH-I3M were present in uninfected plants and accumulated to modest levels at the expense of parent metabolite I3M in flg22- and *Psta*-inoculated plants (Supplementary Fig. 1b) (Clay *et al*., 2009). By comparison, ICN, 4OH-ICN, and camalexin were absent in uninfected plants and at low-to-undetectable levels in plants treated with saturating concentrations of the bacterial MAMPs flg22 and elf26 (10 µM; Felix *et al*., 1999; Zipfel *et al*., 2006). In contrast, ICN, 4OH-ICN and camalexin accumulated to high levels upon inoculation with ETI-inducing pathogens (Fig. 1b; Supplementary Fig. 1c). Furthermore, camalexin, ICN, and 4OH-ICN metabolism was greatly diminished, and indole glucosinolate levels were mostly unchanged in the *rpm1* mutant, which is impaired in ETI recognition of *Psta* (Bisgrove *et al*., 1994) (Supplementary Fig. 1b-c). By contrast, camalexin and ICN were absent in uninfected plants and largely at low-to-undetectable levels in plants treated with MAMPs and PTI/ETS-eliciting pathogens, with 4OH-ICN not detected in most cases. One exception was the fungal MAMP chitosan. 150 µg/mL chitosan induced high levels of camalexin and detectable levels of ICN, consistent with previous observations of camalexin biosynthetic genes upregulation (Fig. 1b) (Povero *et al*., 2011). Higher chitosan concentrations (≥200 µg/mL) have been shown to induce HR-like cell death in Arabidopsis (Cabrera et al., 2006), a phenomenon commonly observed for ETI (Jones and Dangl, 2006). To our surprise, 300 µg/mL chitosan additionally induced detectable levels of 4OH-ICN (Fig. 1b). These results suggest that 4OH-I3M, 4M-I3M, camalexin, and ICN are synthesized in response to multiple PTI elicitors, whereas 4OH-ICN biosynthesis is specific to ETI-like responses.

### *WRKY33* is required to activate 4OH-ICN in response to *Psta*

4OH-ICN biosynthetic genes are highly co-expressed with each other (Rajniak *et al*., 2015) and with camalexin biosynthetic genes (Supplementary Fig. 1d), which are in the WRKY33 regulon (Qiu *et al*., 2008; Birkenbihl *et al*., 2012). To determine whether 4OH-ICN biosynthetic genes are also in the WRKY33 regulon, we compared camalexin, ICN and 4OH-ICN levels between wild-type and a *wrky33* loss-of-function mutant that encodes two differently truncated proteins (Fig. 2a; Zheng *et al*., 2006). Consistent with a previous report (Qiu *et al*., 2008), *wrky33* was impaired in camalexin biosynthesis in response to *Psta* and *Pst avrRps4* (Fig. 2b; Supplementary Fig. 2a). The *wrky33* mutant was similarly impaired in 4OH-ICN biosynthesis (Fig. 2b; Supplementary Fig. 2a). These results indicate that WRKY33 is required for camalexin and 4OH-ICN biosynthesis in response to multiple ETI elicitors.

**Figure 2.**
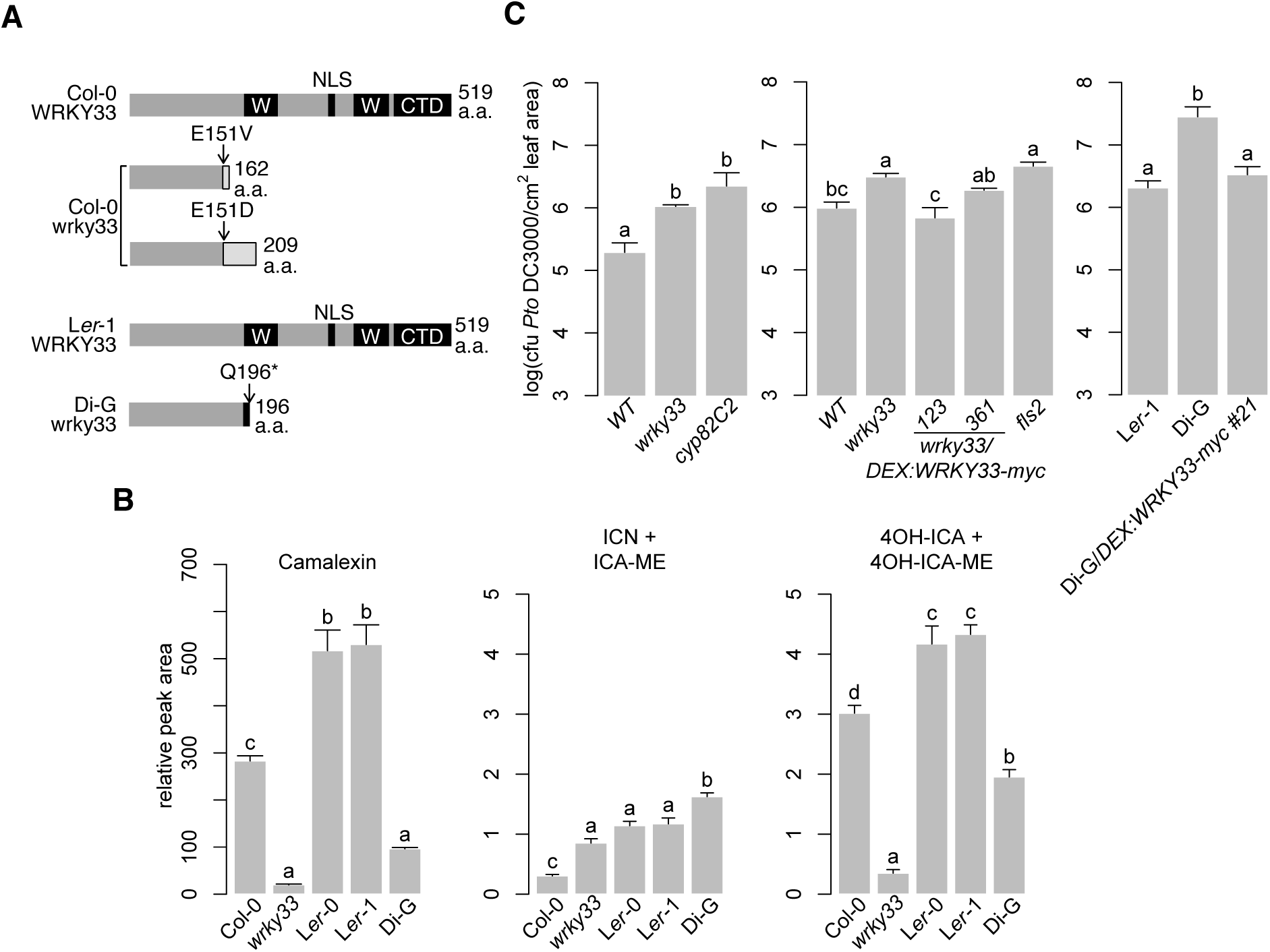
Intraspecific variation in *WRKY33* affects 4OH-ICN and immunity. (**a**) Schematic of WRKY33 proteins in Col-0, Col-0 *wrky33*, L*er*-1 and Di-G. Black boxes denote WRKY domains (W), nuclear localization signal (NLS), or C-terminal domain (CTD). (**b**) LC-DAD-MS analysis of camalexin, ICN, and 4OH-ICN in seedlings inoculated with *Psta* for 24 hr. Data represent mean ± SE of four replicates. (**c**) Bacterial growth analysis of *Pst* in surface-inoculated leaves. Middle and right panels were pre-treated with 20 μM dex for 6-8 hr. Data represent mean ± SE of 4 (left), 6-11 (middle), and 6-8 (right) biological replicates. CFU, colony-forming units. Different letters in (**b-c**) denote statistically significant differences (*P* < 0.05, one-factor ANOVA coupled to Tukey’s test). Experiments in (**b-c**) were performed at least twice, producing similar results. For (**b**-**c**), source data are provided as a Source Data file.

To confirm that *WRKY33* is required to activate the 4OH-ICN pathway, we used a two-component glucocorticoid-inducible system to generate *wrky33* plants that in the presence of the glucocorticoid hormone dexamethasone (dex) express a wild-type copy of WRKY33 with a C-terminal fusion to 1x flag epitope (*wrky33/DEX:WRKY33-flag*; Supplementary Fig. 2b-c). Induced expression of *WRKY33-flag* restored camalexin and 4OH-ICN biosynthesis in *Psta*-challenged *wrky33* plants to greater than wild-type levels (Supplementary Fig. 2d). These results indicate that *WRKY33* is required to activate camalexin and 4OH-ICN biosynthesis in response to *Psta*.

### Intraspecific variation in *WRKY33* affects 4OH-ICN synthesis and pathogen defense

Intraspecific variation in TFs can contribute to gain or loss of phenotypes, such as branching in maize (Studer *et al*., 2011) or pelvic loss in three-spined stickleback fish (Chan *et al*., 2010). In addition, the wide variation in camalexin biosynthesis reported among natural accessions of *A. thaliana* (Kagan & Hammerschmidt, 2002) suggests that a similar variation in 4OH-ICN biosynthesis may exist. To identify additional transcriptional activators of 4OH-ICN biosynthesis that otherwise might be refractory to traditional genetic approaches, we compared intraspecific variation in *Psta*-induced camalexin, ICN and 4OH-ICN among 35 re-sequenced accessions and *wrky33* (Col-0 accession). We found camalexin and 4OH-ICN levels to be positively correlated among accessions (R^2^ = 0.37; Supplementary Fig. 3a), lending further support to their co-regulation by WRKY33. Accession Dijon-G (Di-G) was identified to produce less camalexin and 4OH-ICN and more ICN than its near-isogenic relatives, the Landsberg accessions L*er*-0 and L*er*-1 (Fig. 2b; Supplementary Fig. 3a-b). In addition, differences observed in the metabolite response between Landsberg accessions and Di-G most closely resembled those between Col-0 and *wrky33* mutant (Fig. 2b; Supplementary Fig. 3a). These results led us to hypothesize that genetic variation in a regulatory gene, as opposed to an immune signaling gene, is responsible for the metabolite phenotypes observed in Di-G. To test this hypothesis, genetic variation between Di-G and three sequenced Landsberg accessions (La-0, L*er*-0, and L*er*-1) were used to identify 354 genes that were differentially mutated to high effect in Di-G (Supplementary Fig. 3c). Twenty-eight of these mutated Di-G genes were annotated by Gene Ontology to have roles in defense, including *WRKY3*3 (Supplementary Table 1). We confirmed by Sanger sequencing that Di-G *WRKY33* harbors a nonsense mutation early in the N-terminal DNA-binding motif (Fig. 2a), likely abolishing protein function. Our findings indicate that camalexin and 4OH-ICN are sensitive to intraspecific variation in *WRKY33*.

Camalexin and 4OH-ICN promote plant fitness by contributing non-redundantly to pathogen defense against the fitness-reducing *Pst* (Kover & Scaal, 2002; Rajniak *et al*., 2015). To confirm that disease resistance to *Pst* is also sensitive to intraspecific variation in *WRKY33*, we measured bacterial growth in adult leaves of *wrky33* and Di-G and their respective (near-)isogenic accessions Col-0 and L*er*-1. *wrky33* and Di-G were more susceptible to *Pst* than their (near)isogenic relatives and comparable to the 4OH-ICN biosynthetic mutant *cyp82C2* (Fig. 2c; Rajniak *et al*., 2015).

We additionally generated *wrky33* plants that in the presence of dex express a wild-type copy of WRKY33 with a C-terminal fusion to a larger 6x myc epitope (*wrky33/DEX:WRKY33-myc*; Supplementary Fig. 4a-c). Induced expression of WRKY33-myc complemented *wrky33* and Di-G to Col-0 and L*er*-1 levels of resistance to *Pst*, respectively (Fig. 2c). Additionally camalexin and ICN levels complemented and/or exceeded Col-0 and L*er*-1 levels in *Psta*-challenged *wrky33/DEX:WRKY33-myc* and *Di-G/DEX:WRKY33-myc* plants, respectively (Supplementary Fig. 4d-e). Together, our results support a role of *WRKY33* in pathogen defense as an activator of Trp-derived specialized metabolism.

### WRKY33 activates 4OH-ICN biosynthesis

To confirm that the 4OH-ICN biosynthetic pathway is in the WRKY33 regulon, we first compared *WRKY33*, *CYP71A13, CYP71B15, FOX1* and *CYP82C2* transcript levels among WT, *wrky33*, *wrky33/DEX:WRKY33*-*flag*, and *wrky33/DEX:WRKY33*-*myc*. Consistent with previous reports (Qiu *et al*., 2008), *CYP71A13*, *CYP71B15*, and *FOX1* expression was down-regulated in *wrky33* plants in response to *Psta* and upregulated in both *wrky33/DEX:WRKY33*-*flag* and *wrky33/DEX:WRKY33*-*myc* (Fig. 3a) (Supplementary Fig. 4f, 5a). Interestingly, *CYP82C2* expression and 4OH-ICN production were restored in *wrky33/DEX:WRKY33*-*flag* but not *wrky33/DEX:WRKY33-myc* or Di-G*/DEX:WRKY33-myc* plants (Fig. 2d, 3a) (Supplementary Fig. 4d-f), likely due to the interference of the larger myc tag with the WRKY33 C-terminus, a region previously linked with transactivation activity (Zhou *et al*., 2015). These transcriptional and metabolic findings indicate that WRKY33 mediates camalexin and 4OH-ICN biosynthesis in response to pathogen effectors.

**Figure 3.**
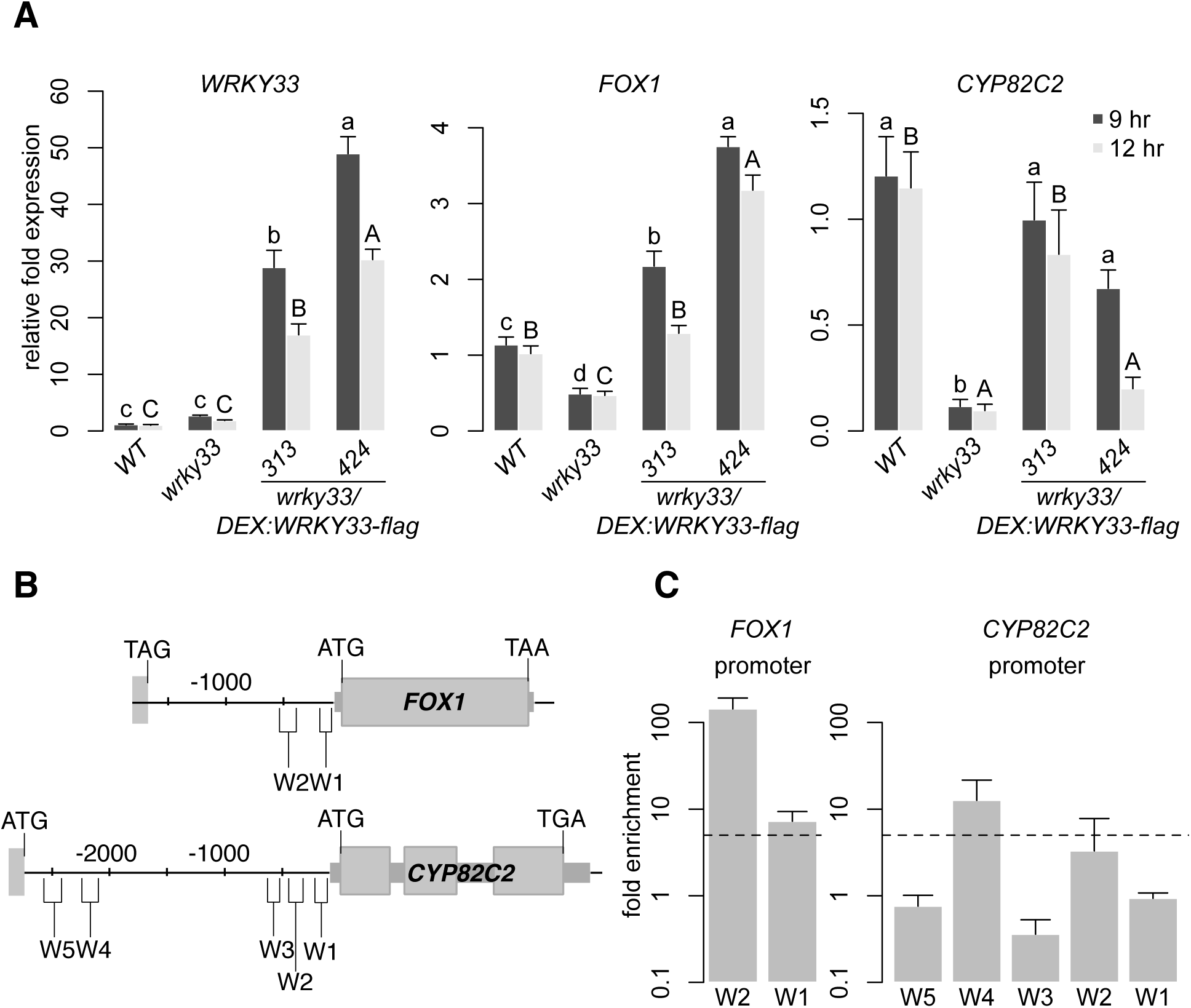
WRKY33 directly activates 4OH-ICN biosynthetic genes. (**a**) qPCR analysis of 4OH-ICN regulatory and biosynthetic genes in seedlings inoculated with 20 µM dex and *Psta* for 9 and 12 hr. Different letters denote statistically significant differences (*P* < 0.05, one-factor ANOVA coupled to Tukey’s test). Lowercase and uppercase letters denote comparisons across 9 and 12 hr timepoints, respectively. Data represent mean ± SE of 4-6 replicates. (**b**) Schematic of *FOX1* and *CYP82C2* loci, indicating nt positions of W-box-containing regions (W). (**c**) ChIP-PCR analysis of W-box-containing regions upstream of *FOX1* and *CYP82C2* in *wrky33/DEX:WRKY33-flag* plants co-treated with 20 µM dex (D) or mock solution (M) and *Psta* for 9 hr. Dashed line represents the 5-fold cutoff between weak and strong TF-DNA interactions. Data represent mean ± SE of four replicates. For (**a**,**c**), source data are provided as a Source Data file.

We then tested for WRKY33-binding to W-box-containing regions upstream of camalexin and 4OH-ICN biosynthetic genes in dex-treated and *Psta*-infected *wrky33/DEX:WRKY33-flag* seedlings by chromatin immunoprecipitation (ChIP)-PCR. WRKY33 has been shown to bind to a W-box region upstream of *CYP71A12* (Birkenbihl *et al*., 2017), a region that also contains three WRKY33-specific motifs and is consistent with WRKY33’s reported binding site preference (Liu *et al*., 2015). We additionally observed that *Psta*-induced WRKY33 bound strongly (greater than 5-fold enrichment) to a single W-box region upstream of *FOX1* and *CYP82C2* (W2 and W4, respectively; Fig. 3b-c; Supplementary Fig. 5b). Both regions also contain one to three WRKY33-specific motifs. Together with our expression analysis, our findings indicate that WRKY33 uses preferred WRKY33-binding sites to directly activate 4OH-ICN biosynthetic genes in response to pathogen effectors.

Interestingly, *Psta*-induced WRKY33 did not bind to the W5 region upstream of *CYP82C2* (Fig. 3c), a W-box region that does not contain any WRKY33-specific motifs and is just upstream of neighboring gene of unknown function *At4g31960* (Fig. 3b). WRKY33 reportedly binds to W5 in response to flg22 and *B. cinerea* (Liu *et al*., 2015; Birkenbihl *et al*., 2017). By contrast, *Psta*-induced WRKY33 bound strongly to W1 region upstream of *CYP71B15* (Supplementary Fig. 5c-d), a W-box region that also does not contain any WRKY33-specific motifs. WRKY33 reportedly binds to a region encompassing W1 in response to flg22 and *Psta* (Qiu *et al*., 2008; Birkenbihl *et al*., 2012). These findings suggest that WRKY33 may use W-box extended motifs or novel specificity motifs to target camalexin biosynthetic genes in response to pathogen effectors, or 4OH-ICN biosynthetic genes in response to MAMPs or fungal pathogens.

### *CYP82C2* underwent regulatory neofunctionalization

CYP82C2 catalyzes the last step in 4OH-ICN biosynthesis, hydroxylating ICN to form 4OH-ICN (Rajniak *et al*., 2015), and likely was the last 4OH-ICN pathway gene to be recruited to the WRKY33 regulon in *A. thaliana*. To explore the phylogenetic distribution pattern of 4OH-ICN biosynthesis, we profiled ICN and 4OH-ICN metabolites in close and distant relatives of *A. thaliana* in response to *Psta*. While ICN biosynthesis was observed across multiple close relatives, 4OH-ICN was only detected in *A. thaliana* (Fig. 4a; Supplementary Fig. 6a). This result suggests that 4OH-ICN manifests a species-specific diversification of pathogen-inducible Trp-derived metabolism in the mustard family.

**Figure 4.**
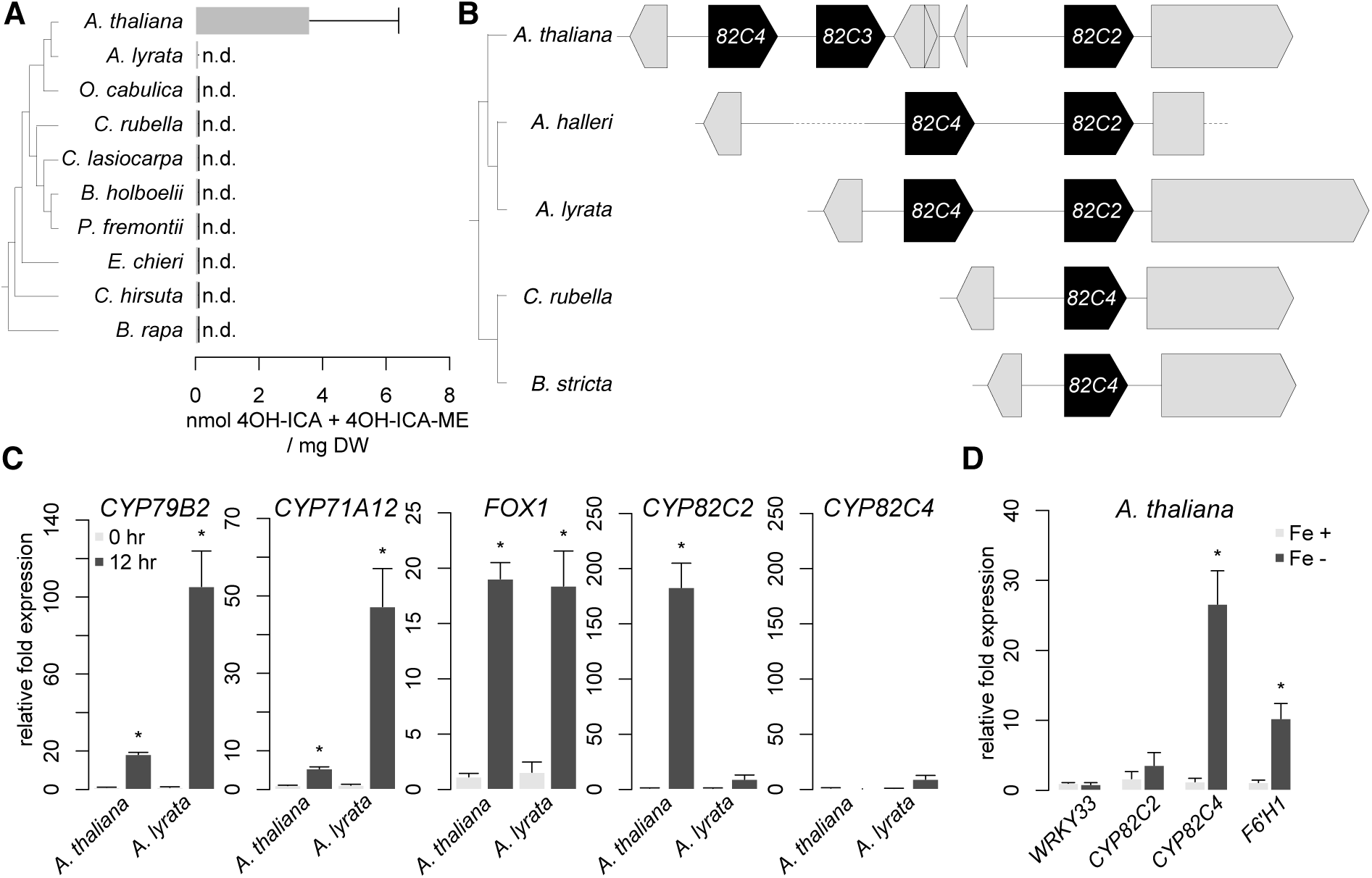
Regulatory neofunctionalization of *CYP82C2*. (**a**) (Left) Phylogenetic species tree. (Right) HPLC-DAD analysis of 4OH-ICN in seedlings inoculated with *Psta* for 30 hr. Data represent mean ± SE of three independent experiments (n = 4 biological replicates), each with *A. thaliana* as a positive control. 4OH-ICA and 4OH-ICA-ME are aqueous and methanolic degradations products of 4OH-ICN, respectively. DW, dry weight; n.d., not detected. (**b**) (Left) phylogenetic species tree. (Right) Synteny map of the *CYP82C* genes. Grey arrows or rectangles represent non-*CYP82C* genes. Grey dotted lines represent large (>500 nt) sequence gaps. (**c-d**) qPCR analysis of 4OH-ICN and sideretin biosynthetic genes in seedlings inoculated with *Psta* (**c**) or grown in iron-deficient medium (**d**). Data represents the mean ± SE of four biological replicates.Asterisks denote statistically significant differences of stress-treated relative to untreated samples (*P* < 0.05, two-tailed t-test). For (**a**,**c-d**), source data are provided as a Source Data file.

In *A. thaliana*, *CYP82C2* resides in a near-tandem cluster with paralogs *CYP82C3* and *CYP82C4* (Fig. 4b). We performed phylogenetic and syntenic analyses to identify putative *CYP82C2* orthologs in ICN-synthesizing species. All identified homologs are syntenic to *CYP82C2* or *CYP82C4*, and encode proteins with >88% identity to one another (Fig. 4b; Supplementary Fig. 6b-c). *CYP82C3* is present only in *A. thaliana*, and although more similar to *CYP82C2* than *CYP82C4* in sequence, it is not functionally redundant with *CYP82C2* (Fig. 4b; Supplementary Fig. 6b; Rajniak *et al*., 2015). *CYP82C4* is required for the biosynthesis of sideretin, a widely conserved, phenylalanine-derived metabolite required for iron acquisition (Rajniak *et al*., 2018). *CYP82C4* has syntenic orthologs in the mustard family, correlating with the distribution of sideretin biosynthesis (Fig. 4b; Supplementary Fig. 6b; Rajniak *et al*., 2018). By contrast, *CYP82C2* has syntenic orthologs only within the *Arabidopsis* genus (Fig. 4b; Supplementary Fig. 6b). These results suggest that *CYP82C2* duplicated from *CYP82C4* prior to the formation of the *Arabidopsis* genus and then acquired a new expression pattern and/or catalytic function prior to *A. thaliana* speciation approx. 2 million years later (Hu *et al*., 2011; Hohmann *et al*., 2015).

CYP82C2 and CYP82C4 were previously characterized to 5-hydroxylate with equal efficiency the specialized metabolite 8-methoxypsoralen, a molecule structurally reminiscent of ICN and sideretin (Kruse *et al*., 2008). The apparent similarities in substrate specificity and catalytic function suggest that *CYP82C2* may have diverged from *CYP82C4* in expression but not protein function. To test this, we first compared the expression of *CYP82C2* and *CYP82C4* in *A. lyrata* and *A. thaliana* in response to *Psta*. 4OH-ICN biosynthetic genes *CYP79B2, CYP71A12* and *FOX1* were upregulated in both species, consistent with the common presence of ICN (Fig. 4a and c). By contrast, *CYP82C2* levels were respectively upregulated and unchanged in *A. thaliana* and *A. lyrata*, correlating with the distribution of 4OH-ICN in these species (Fig. 4a and c). *CYP82C4* expression was unchanged in both species (Fig. 4c). These results indicate that 4OH-ICN biosynthesis is linked with pathogen-induced expression of *CYP82C2*.

We then compared the aligned upstream sequences of *CYP82C2* and *CYP82C4* in *A. lyrata* and *A. thaliana* and observed good sequence conservation among orthologs but poor conservation among paralogs (Supplementary Fig. 6d), indicating that sequences upstream of *CYP82C4* and *CYP82C2* were independently derived. We performed expression analysis in *A. thaliana* to confirm that *CYP82C2* and *CYP82C4* have different expression patterns. *CYP82C2* expression is upregulated in response to *Psta* and unchanged under iron deficiency (Fig. 4c-d; Supplementary Fig. 1a; Rajniak *et al*., 2015). Conversely, *CYP82C4* is upregulated under iron deficiency and unchanged in response to *Psta* (Fig. 4c-d; Murgia *et al*., 2011; Rajniak *et al*., 2018). Finally, *CYP82C4* was unchanged in *Psta*-challenged *wrky33* and *wrky33/DEX:WRKY33-flag* (Supplementary Fig. 6e). Our findings suggest that *CYP82C2* diverged from *CYP82C4* by acquiring WRKY33 regulation for its pathogen-induced expression.

We next assessed dN/dS ratios along branches of the CYP82C phylogenetic tree (Supplementary Fig. 6b) and found good support for purifying selection acting on CYP82C enzymes (ω=0.21), and no support for positive selection acting on CYP82C2/3 enzymes (Supplementary Table 2). Lastly, we identified non-conserved amino acid residues among CYP82C homologs and mapped this information onto a homology model of CYP82C2. The protein inner core, which encompasses the active site and substrate channel, is highly conserved among CYP82C homologs (Supplementary Fig. 6f), and is consistent with CYP82C2 and CYP82C4’s reportedly redundant catalytic functions (Kruse *et al*., 2008). Altogether, our findings suggest that *CYP82C2* underwent regulatory neofunctionalization (Moore & Purugganan, 2005), diverging from *CYP82C4* in expression but not protein function.

### TE *EPCOT3* is a *CYP82C2* enhancer

WRKY33 regulation of *CYP82C2* is mediated by a WRKY33-TFBS in the W4 region (Figs. 3 and 5a; Supplementary Fig. 5c). Preferential WRKY33-binding at this region should also be influenced by chromatin features associated with *cis*-regulatory elements like enhancers and basal promoters (Slattery *et al*., 2014). To investigate how *CYP82C2* acquired WRKY33-binding for its pathogen-induced expression, we compared the aligned upstream sequences of *CYP82C* homologs in ICN-synthesizing species. We observed three large upstream sequences specific to *A. thaliana CYP82C2*, hereafter named ***<u>E</u>****ighty-two-C2 **<u>P</u>**romoter **<u>C</u>**ontained **<u>O</u>**nly in A. **<u>T</u>**haliana1-3* (*EPCOT1–3*; Fig. 5a). *EPCOT3* in particular is a 240nt region that completely encompasses W4 (Fig. 5a), indicating that the WRKY33’s regulation of *CYP82C2* in response to *Psta* may be species-specific. Further bioinformatics analysis revealed that *EPCOT3* has the epigenetic signature of an active enhancer (Roudier *et al*., 2011; Liu *et al*., 2018). Relative to neighboring sequences, *EPCOT3* is enriched with activating histone mark H3K4me2 and lacks the repressive histone mark H3K27me3 (Fig. 5b) (Heintzman *et al*., 2007; Hoffman *et al*., 2010; Roudier *et al*., 2011; Bonn *et al*., 2012; Wang *et al*., 2014). Our findings suggest that *EPCOT3* functions as an enhancer that mediates WRKY33-binding and activation of *CYP82C2* in response to pathogen effectors.

**Figure 5.**
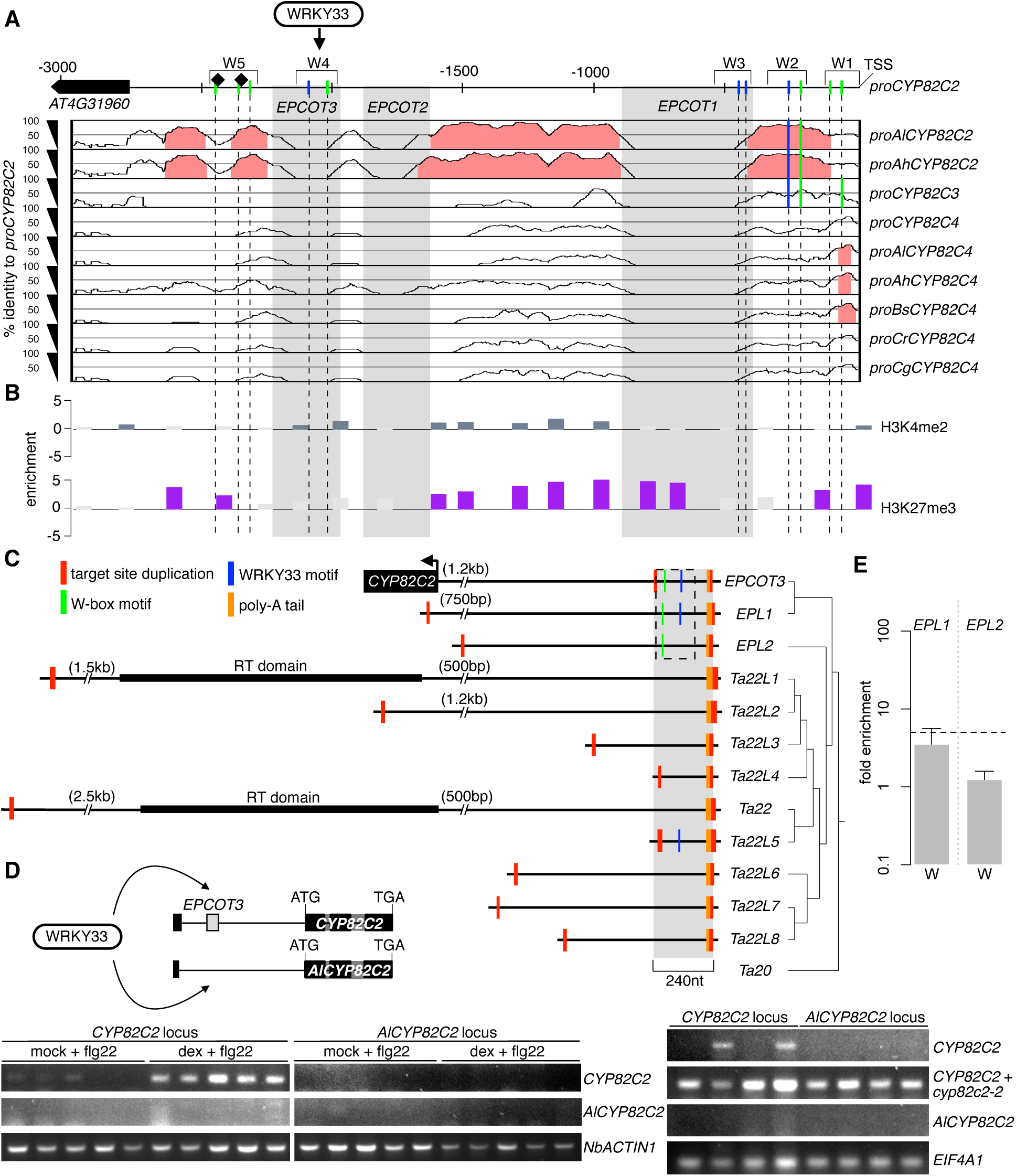
TE *EPCOT3* is a *CYP82C2* enhancer. (**a**) mVISTA plot of *CYP82C2* upstream sequence, indicating nt positions of unique (*EPCOT1–3;* gray boxes) and conserved regions (≥70% sequence identity; pink) among homologous sequences. Also indicated are positions of W-boxes (green) and WRKY33-specific motifs (blue) that are present (solid lines) or absent (dashed lines) in each homologous sequence, previously known WRKY33-TFBSs (diamonds) and ChIP-tested regions (W1-5). TSS, transciptional start site; *Al, Arabidopsis lyrata; Ah, Arabidopsis halleri*; *Cr, Capsella rubella*; *Bs, Boechera stricta; Cg, Capsella grandiflora*. (**b**) Epigenetic map of *CYP82C2* upstream sequence, indicating nt positions of significant amounts of H3K4me2 (blue-gray bars), and H3K27me3 (purple bars). (**c**) (Left) Schematic of *EPCOT3* and related LINE retrotransposons in *A. thaliana* drawn to scale, indicating nt positions of *CYP82C2* and reverse transcriptase (RT) domain. A more detailed tree is available as Supplementary Text 1. (Right) Phylogenetic maximum likelihood tree. Dashed box represent region containing W-boxes (green lines) and/or WRKY33-binding motifs (blue lines) within *EPCOT3, EPL1* and *EPL2*. (**d**) (Upper left) Schematic of *CYP82C2* and *AlCYP82C2* transgenic loci used for WRKY33 transactivation experiments. Black box at far left represents coding sequences of neighboring genes (*AT4G31960* at *CYP82C2* locus, *AlCYP82C4* at *AlCYP82C2* locus). (Lower left) RT-PCR images of *CYP82C2*, *AlCYP82C2*, and *NbACTIN1* in *N. benthamiana* leaves co-transfected with *DEX:WRKY33-flag* and the *CYP82C2* or *AlCYP82C2* locus and incubated with 1 μM flg22 and either mock solution (0.5% DMSO) or 20 μM dex for 30 hr (*CYP82C2* and *AlCYP82C2*) or 24 hr (*NbACTIN1*). Data represents five biological replicates. (Lower right) RT-PCR images of *CYP82C2*, *AlCYP82C2*, and *EIF4A1* in *A. thaliana cyp82C2-2* mutant protoplasts transfected with the wild-type *CYP82C2* or *AlCYP82C2* locus and elicited with 1 μM flg22 for 6 hr. As transcription is present downstream of the T-DNA insertion in the *cyp82C2-2* mutant, primers for RT-PCR were selected corresponding to a downstream region (*CYP82C2* + *cyp82c2-2*, second row) as well as a region flanking the *cyp82C2-2* insertion (*CYP82C2*, first row). See Supplementary Table 6 for more information. Data represents four biological replicates. (**e**) ChIP-PCR analysis of W-box-containing regions (W) within *EPL1* and *EPL2* in *wrky33/DEX:WRKY33-flag* plants co-treated with 20 µM dex (D) or mock solution (M) and *Psta* for 9 hr. Data represent mean ± SE of four replicates. Dashed line represents the 5-fold cutoff between weak and strong TF-DNA interactions. For (**d-e**), source data are provided as a Source Data file.

*EPCOT3* contains a 3’ poly-A tail and is flanked by variable-length target site duplications (Fig. 5c; Supplementary Fig. 7a), which are hallmarks of eukaryotic LINE retrotransposons (Malik *et al*., 1999). LINE retrotransposition (reverse transcription and integration) results in frequent 5’-truncation of retrocopies (Luan *et al*., 1993). We identified eleven variably truncated retrocopies similar to *EPCOT3* throughout the genome, including *Ta22*, one of the first LINEs characterized in *A. thaliana* (Fig. 5c; Supplementary Fig. 7a-b, Supplementary Table 3; Wright *et al*., 1996). *EPCOT3*-related LINEs were sorted into two groups roughly correspondent to their phylogenetic placement: *EPCOT3-LIKE* (*EPL*) for those with high identity (>65%) to *EPCOT3* and *Ta22* or *Ta22*-*LIKE* (*Ta22L*) for the remainder (Supplementary Fig. 7a; Supplementary Table 3). Only *Ta22* and *Ta22L1* are full-length LINEs (Fig. 5c), presumably encoding the proteins necessary for their own transposition and for the transposition of nonautonomous family members like *EPCOT3*. We also identified two syntenic species-specific *Ta22L*s, but no *EPLs*, in *A. lyrata* (Supplementary Table 3). To confirm the involvement of *EPCOT3* in species-specific expression of *CYP82C2*, we introduced WRKY33 into *N. benthamiana* leaves and *A. thaliana cyp82C2* protoplasts transfected with either the *A. thaliana* and *A. lyrata CYP82C2* loci (including genes plus 3000 bp upstream sequence, Fig 5d). We observed transactivation by WRKY33 of the *A. thaliana* sequence, but not that of *A. lyrata* (Fig. 5d; Supplementary Fig. 7d). Altogether, these data indicate that *EPCOT3* and *EPLs* arose from retrotransposition following the speciation of *A. thaliana* and that the *EPCOT3*-containing *A. thaliana CYP82C2* promoter is sufficient to confer WRKY33-mediated transcription of *CYP82C2*. Of the *EPL* retrocopies, *EPL1* is most similar to *EPCOT3* (85.4% identity), sharing the W-box and WRKY33-specific motif, whereas *EPL2* is less similar (67%) and lacks the WRKY33-specific motif (Fig. 5c; Supplementary Table 3, Supplementary Fig. 7a). *EPL1* and *EPL2* are much less truncated than *EPCOT3* (Fig. 5c), and lack epigenetic signatures typical of *cis*-regulatory sequences (Supplementary Fig. 7c) (Roudier *et al*., 2011; Liu *et al*., 2018). To investigate whether the sequence information and chromatin features associated with *EPL*s are sufficient for WRKY33-binding, we tested for WRKY33-binding to *EPL* sequences homologous to the W4 region of *EPCOT3* in dex-treated, *Psta*-infected *wrky33/DEX:WRKY33-flag* plants by ChIP-(q)PCR. Compared to *EPCOT3* (Fig. 3c), WRKY33 respectively bound weakly or not at all to *EPL1* and *EPL2* (Fig. 5e; Supplementary Fig. 7e). Our findings suggest the following history: (1) *EPL1* likely retroduplicated from *EPL2* or its progenitor, which already contained a W-box; (2) *EPL1* then acquired a WRKY33-specific motif by mutation; (3) *EPCOT3* retroduplicated from *EPL1* and then acquired epigenetic signatures of an enhancer, thereby allowing selection to act on standing variation rather than *de novo* mutation for *CYP82C2* recruitment into the 4OH-ICN biosynthetic pathway.

## Discussion

TEs were originally conceived to act as “controlling elements” of several loci in the genome (McClintock, 1956), and exaptation of TEs into *cis*-regulatory modules has been hypothesized to be responsible for the rapid transcriptional rewiring in more ancient lineages of vertebrates (Feschotte 2008; Bourque 2009; de Souza et al., 2013). However, few (if any) evolutionarily recent TE exaptation events in vertebrates and higher plants have been demonstrated to have biochemical, regulatory, physiological and fitness-promoting functions (de Souza et al., 2013). With well over a dozen genomes available including the genetic model *A. thaliana*, the mustard family presents an excellent system for examining such events. In this study, we show that *EPCOT3* is a TE-derived enhancer that mediates WRKY33-binding, pathogen-responsive transcription of *CYP82C2,* synthesis of the species-specific metabolite 4OH-ICN, and pathogen defense (Fig. 6). These results provide the first instance of a recent TE exaptation responsible for the rewiring of a new gene into an ancient regulon, ultimately leading to a positive effect on fitness.

**Figure 6.**
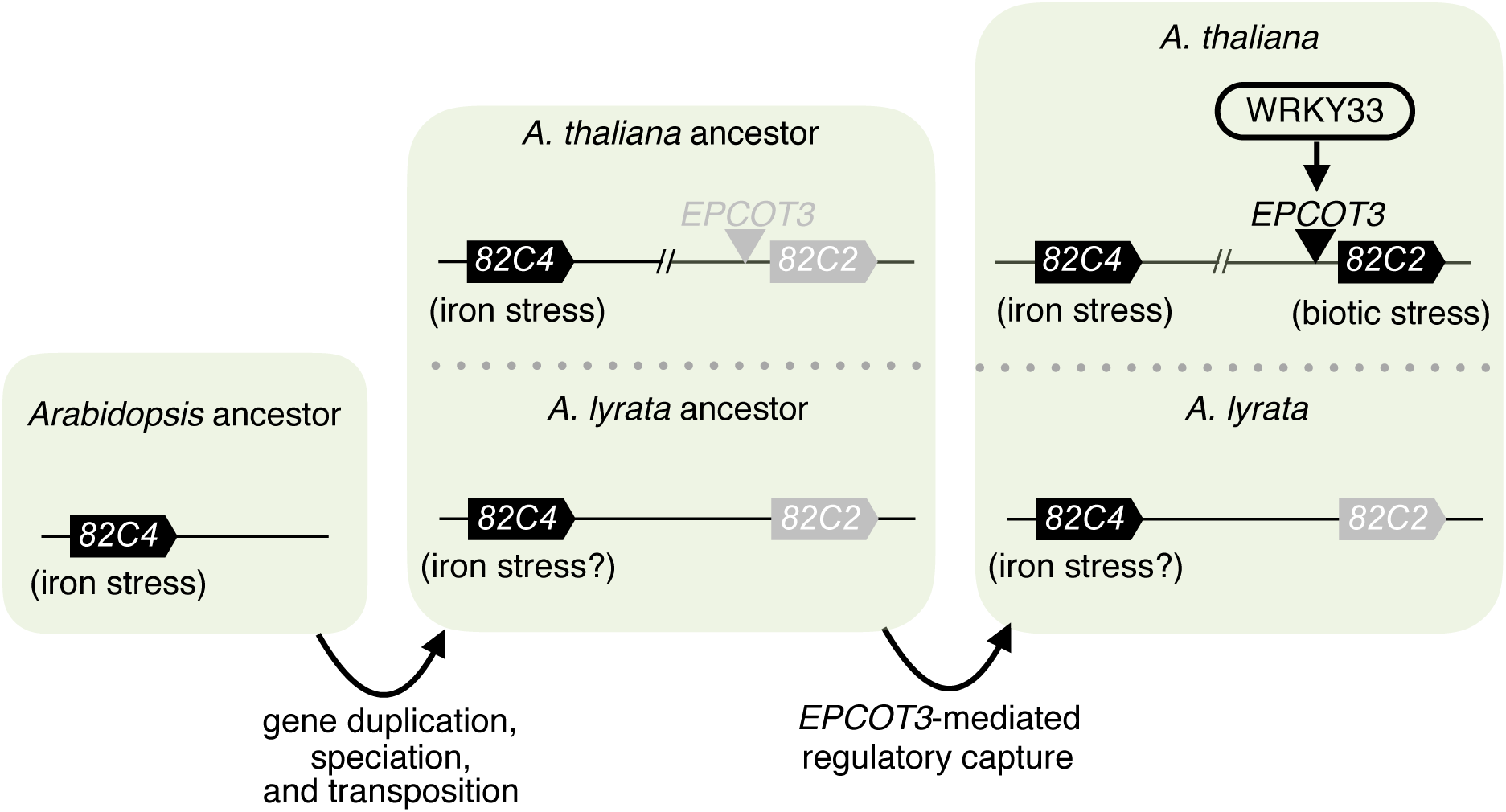
Model of regulatory neofunctionalization of *CYP82C2*.

Although the *EPL1/EPCOT3* progenitor retrotransposed a preferred WRKY33-TFBS in the form of *EPCOT3* upstream of *CYP82C2*, a further series of epigenetic modifications were needed to facilitate optimal access of *EPCOT3* by WRKY33 (Fig. 6). *EPL1* exists in a silenced heterochromatin state (Supplementary Fig. 7c), typical for TEs (Slotkin & Martienssen, 2007), and is bound weakly by WRKY33 (Fig. 5e), whereas *EPCOT3* is in an open chromatin state (Fig. 5b; Roudier *et al*., 2011; Liu *et al*., 2018) and bound strongly by WRKY33 (Fig. 3c). The more severe 5’-truncation of *EPCOT3* could account for its release from TE silencing mechanisms, and the initially weak WRKY33-binding could provide a ‘seed’ for chromatin remodelers to drive the exaptation of newly retrotransposed *EPCOT3* into a *bona fide* enhancer. Further epigenomic sampling within *Arabidopsis* is needed to better clarify the epigenetic transformations underlying the *EPCOT3* exaptation event.

Compared to closely-related Landsberg accessions (Supplementary Fig. 3; Hardtke *et al*., 1996), Di-G synthesizes less camalexin and 4OH-ICN (Fig. 2b; Kagan & Hammerschmidt, 2002), is more susceptible to a range of bacterial and fungal pathogens (Fig. 2c) (Hugouvieux *et al*., 1998; Kagan & Hammerschmidt, 2002; Mukherjee *et al*., 2009), and is more sensitive to the phytohormone ethylene (Chatfield *et al*., 2008). WRKY33 has been implicated in camalexin biosynthesis (Qiu *et al*., 2008), antifungal defense (Zheng *et al*., 2006), and ethylene biosynthesis (Li *et al*., 2012). We identified WRKY33 as causal for some if not all of these phenotypes in Di-G. This is the first report of WRKY33’s involvement in antibacterial defense and is consistent with the contribution of camalexin and 4OH-ICN towards antibacterial defense (Rajniak *et al*., 2015).

WRKY33 is an ancient transcription factor responsible for many fitness-promoting traits in plants, thus it is unexpected that an *A. thaliana* accession would have a naturally occurring *wrky33* mutation (C536T transversion). Di-G is the sole member of 1,135 sequenced accessions to have a high-effect single nucleotide polymorphism (SNP) in *WRKY33* (1001 Genomes Consortium, 2016). Di-G and L*er*-0 have long been models for studies in mutagenesis (Rédei, 1962, Müller, 1966), and thus a possibility exists that Di-G may have originated from an ethyl methanesulfonate (EMS) mutagenesis screen of L*er*-0. Historical EMS mutagenesis experiments generated upwards of tens of thousands of mutations per cell (Müller 1966; Rédei & Koncz, 1993; Camara *et al*., 2000), well within the range of ∼25,000 SNPs that are not concordant between Di-G and L*er*-0 (Supplementary Fig. 2f). However, features of EMS mutations (i.e. transversion mutations) or X-ray mutations (i.e. indels) are not enriched in the Di-G pseudogenome relative to related pseudogenomes (Supplementary Table 4). These findings suggest that the wrky33 Di-G mutation is naturally derived.

## Methods

### Plant materials and growth conditions

For qPCR and HPLC-DAD analyses, surface-sterilized *Arabidopsis thaliana* seeds were sown in 12-well microtiter plates sealed with Micropore tape (3M, St. Paul, MN), each well containing ∼15 seeds and 1 mL filter-sterilized 1X Murashige and Skoog (MS; Murashige & Skoog, 1962) media (pH 5.7–5.8) (4.43 g/L MS basal medium with vitamins [Phytotechnology Laboratories, Shawnee Missions, KS], 0.05% MES hydrate, 0.5% sucrose). Iron deficient media was made as previously described by Rajniak et al (2018). For *Polyctenium fremontii*, surface-sterilized seeds were sown on MS agar plates. On day 9, seedlings were transferred to 6-well microtiter plates, each well containing ∼15 seeds and 3 mL MS media. For all other species, surface-sterilized seeds were sown in 6-well plates, each well containing ∼15 seeds and 3 mL MS media. On day 9, media were refreshed prior to bacterial elicitation. Microtiter plates were placed on grid-like shelves over water trays on a Floralight cart (Toronto, Canada), and plants were grown at 21°C with 60% humidity under a 16-hr light cycle (70-80 μE m-2 s-1 light intensity). For chromatin immunoprecipitation analyses, approximately 200 seeds were sown in a 100mm x 15mm petri plate containing 20mL of 1X MS media. Media were exchanged for fresh media on day 9. Microtiter plates were placed on grid-like shelves over water trays on a Floralight cart (Toronto, Canada), and plants were grown at 21°C with 60% humidity under a 16-hr light cycle (70-80 μE m-2 s-1 light intensity). For bacterial infection assays, seedlings were transferred to and grown on soil [3:1 mix of Farfard Growing Mix 2 (Sun Gro Horticulture, Vancouver, Canada) to vermiculite (Scotts, Marysville, OH)] at 22°C daytime/18°C nighttime with 60% humidity under a 12-hr light cycle [50 (dawn/dusk) and 100 (midday) μE m-2 s-1 light intensity]. Seed stock information is shown in Supplementary Table 5.

### Vector construction and transformation

To generate the *DEX:WRKY33-flag* construct, *WRKY33* was PCR-amplified from genomic DNA using the primers WRKY33gXhoF (5’-AACTCGAGAAGAACAAGAACCATCAC-3’), and W33flgSpeIR (5’-CGACTAGTCTACTTGTCGTCATCGTCTTTGTAGTCGGGCATAAACGAATCGAAA-3’) and subcloned into the *Xho*I/*Spe*I sites of pTA7002 vector (Aoyama and Chua, 1997; McNellis et al., 1998). To generate the *DEX:WRKY33-myc* construct, *WRKY33* was PCR-amplified using the primers WRKY33gXhoF and WRKY33gStuR (5’-AAGGCCTGGCATAAACGAATCGAAAAATG-3’) and subcloned into the *Xho*I/*Stu*I sites of a version of pTA7002 modified to contain 6 tandem copies of the c-Myc epitope downstream of the StuI site (Chezem *et al*., 2017). The constructs were introduced into *Arabidopsis thaliana wrky33* plants via *Agrobacterium*-mediated floral dip method (Clough and Bent, 1998), and transformants were selected on agar media containing 15 μg/mL hygromycin B (Invitrogen, Carlsbad, CA). To generate the *CYP82C2pro:CYP82C2* DNA construct, the *CYP82C2* promoter (pro) and coding sequences were PCR-amplified from *A. thaliana* genomic DNA using the primers At82C2proXbaF (5-GCTCTAGAAGCTTCCAATAAAACATTC-3’) and At82C2proBamR (5’-GCGGATCCAGTGGTTTGAGCGTGCAAA-3’), and At82C2geneBamF (5-GCGGATCCATGGATACTTCCCTCTTTTC-3’) and At82C2geneSmaR (5’-TTCCCGGGCTACTTGTCGTCATCGTCTTTGTAGTCCACATAAAGCCCTTCCTTAAG-3’), and both sequences were subcloned into the *XbaI/SmaI* sites of pBI101 vector (Jefferson *et al.,* 1987). To generate the *AlCYP82C2pro:AlCYP82C2* DNA construct, the *AlCYP82C2* pro and coding sequences were PCR-amplified from *A.lyrata* genomic DNA using the primers Al82C2proSalF (5’-CGGTCGACTATTCCAGGAGCATACAA-3’) and Al82C2proBglIIR (5’-GGAGATCTAATGTTTTAAAAGTGCAAAAGAG-3’), and Al82C2geneBamHF (5’-GCGGATCCATGGATACATCCCTCTTTTC-3’) and Al82C2geneSmaR (5’-TTCCCGGGctacttgtcgtcatcgtctttgtagtccacaaaaagttcttccttaagac-3’), and subcloned into the *SalI/SmaI* sites of pBI101 vector. Transient expression of constructs in *Nicotiana benthamiana* leaves was performed as previously described (Rajniak *et al.,* 2015) with the following modification: leaves were infiltrated with transformed *Agrobacterium* strains that were grown in LB medium supplemented with 30 µg/mL gentamycin and 50 µg/mL kanamycin to an OD_600_ of 0.7. 16 h post-*Agro*-infiltration, leaves were sprayed with 20 µM dexamethasone and 0.1% Tween-20 to induce transgene expression for 0.5, 15, 24 and 30 h. Transient expression of constructs in *Arabidopsis thaliana* T-DNA insertion mutant *cyp82C2-2* (Arabidopsis Biological Resource Center; GABI_261D12) was performed as previously described (Sheen, 2002) with the following modification: 2.5 x 10^5^ protoplasts were transfected with 3 µg of construct for 20 min, recovered in 2.5x volume of W5 solution and elicited with 1 µM flg22 (QRLSTGSRINSAKDDAAGLQIA; Genscript, Nanjing, China) in 1 mL W5 solution for 6 h.

### Bacterial infection and MAMP elicitation

A single colony of *Pseudomonas syringae* pv. *maculicola* (*Pma*) M2 (containing *avrRpm1*, but not *avrRps4* or *avrRpt2*), *Pma* ES4326 (containing no aforementioned effectors), *Pma* ES4326 *avrRpt2*, *Pseudomonas syringae* pv. *tomato* DC3000 (*Pto* DC3000 or *Pst,* containing no aforementioned effectors), *Pst avrRpm1*, *Pst avrRps4*, and *Pst avrRpt2* from a freshly streaked 3-day-old agar plate were used to inoculate 2 mL of LB containing appropriate antibiotics. Strains were grown to log phase, washed in sterile water twice, resuspended in water to OD_600_ of 0.2, and incubated at room temperature with no agitation for 3-6 and prior to infection. Chitosan (90% deacetylated chitin; Spectrum Chemical Mfg Corp, New Brunswich, NJ) was prepared in 0.1 N acetic acid and neutralized with 0.1 N NaOH to pH 5.8 to a stock concentration of 1.2 mg/mL.

Hydroponically grown 9-day-old seedlings were inoculated with bacterial strains to OD_600_ of 0.013 or treated with 10 μM flg22, 10 μM elf26 (ac-SKEKFERTKPHVNVGTIGHVDHGKTT; Genscript), and 150 or 300 μg/mL chitosan.

For qPCR analyses, seedlings were snap-frozen in liquid nitrogen 12 hr post-infection. For HPLC-DAD analyses, seedlings were snap-frozen 24 to 28 hr post-infection. For ChIP analyses, seedlings were snap-frozen 9 hr post-infection.

4-to-5-week-old adult leaves were treated with 0.0125% Silwet or 0.0125% Silwet and 20 μM dexamethasone for 20 sec and incubated on 0.8% (w/v) tissue-culture agar plates on a light cart at 21°C for 6-8 hr. Leaves were then surface-inoculated with *Pto* DC3000 (OD_600_ = 0.002 or 10^6^ colony-forming units (cfu)/cm^2^ leaf area) in the presence of 0.01% (v/v) Silwet L-77 (Phytotechnology Laboratories) for 15 sec and incubated on 0.8% (w/v) tissue-culture agar plates at 21°C under a 16-hr light cycle (70-80 μE m-2 s-1 light intensity) for 3 days. Leaves were then surface-sterilized in 70% ethanol for 10 sec, rinsed in sterile water, surface-dried on paper towels, and the bacteria were extracted into water, using an 8-mm stainless steel bead and a ball mill (20 Hz for 3 min). Serial dilutions of the extracted bacteria were plated on LB agar plates for colony-forming units (CFU) counting.

### RNA isolation and quantitative PCR (qPCR)

Total RNA was extracted from 9-day-old seedlings using TRIzol reagent (Invitrogen, San Diego, CA) according to the manufacturer’s instructions. 2.5 μg of total RNA was reverse-transcribed with 3.75 μM random hexamers (Qiagen, Hilden, Germany) and 20 U of ProtoScript II (New England Biolabs, Boston, MA) according to the manufacturer’s instructions. The resulting cDNA:RNA hybrids were treated with 10 U of DNase I (Roche, Basel, Switzerland) for 30 min at 37°C and purified on PCR clean-up column (Macherey-Nagel, Düren, Germany). qPCR was performed with Kapa SYBR Fast qPCR master mix (Kapa Biosystems, Wilmington, MA) and CFX384 real-time PCR machine (Bio-Rad, Hercules, CA). The thermal cycling program was as follows: 95°C for 3 min; 45 cycles of 95°C for 15 sec and 53°C for 30 sec; a cycle of 95°C for 1 min, 53°C for 1 min, and 70°C for 10 sec; and 50 cycles of 0.5°C increments for 10 sec. Biological and technical replicates were performed on the same 384-well PCR plate. Average of the three Ct values per biological replicate was converted to difference in Ct value relative to that of control sample. The Pfaffl method (Pfaffl, 2001) and calculated primer efficiencies were used to determine the relative fold increase of the target gene transcript over the *EIF4A1* (*AT3G13920* or *AL3G26100*) housekeeping gene transcript for each biological replicate. Expression values were then calculated relative to WT un-treated samples. Primer sequences and efficiencies are listed in Supplementary Table 6.

### RT-PCR Analysis

Total RNA was extracted from snap-frozen tobacco leaves or Arabidopsis protoplasts as described by Couto et al (2015) and Oñate-Sánchez and Vicente-Carbajosa (2008). Total RNA (2 µg) was treated with 10 units of DNase I (Roche) for 30 min at 37°C and then for 15 min at 70°C, reverse-transcribed with 3.75 µM random hexamers (Qiagen) and 20 units of M-MuLV (New England Biolabs) for 1 h at 42°C and then for 15 min at 70°C. Tobacco cDNA was diluted 7.5-fold. 4 µL of cDNA was used in 20 µL PCR reactions, and resulting PCR products were separated on 2% agarose gels. PCR was performed on C1000 thermal cycler (Bio-Rad) with the following thermal cycling program: 95°C for 3 min; 40 cycles of 95°C for 10 sec, 53°C for 15 sec, and 72°C for 7 sec (*AtWRKY33*, *CYP82C2*, *AlCYP82C2*), 15 sec (*NbACTIN1*) or 21 sec (*CYP82C2*-long). Primer sequences are listed in Supplementary Table 6.

### Camalexin and 4OH-ICN extraction and LC-DAD-MS

10-day-old seedlings were snap-frozen, lyophilized, weighed and homogenized using a 5-mm stainless steel bead and ball mill (20 Hz, 4 min). For phytoalexin analysis, homogenate was extracted with 300 μL 80% (v/v) aqueous methanol containing 0.08% (v/v) formate and 2.5 μL internal

standard (200 μM 4-methoxyindole/4M-I [Sigma-Aldrich] in 100% methanol) per mg sample dry weight. Extracts were sonicated for 5 min and centrifuged at 16,000xg for 2 min. The supernatant was filtered using a 0.45-μm polypropylene filter plate (GE Healthcare, Chicago, IL). Samples were separated by reversed-phase chromatography on an Ultimate 3000 HPLC (Dionex, Sunnyvale, CA) system, using a 3.5-μm, 3x150-mm Zorbax SB-Aq column (Agilent, Santa Clara, CA); volume injected was 10 μL. The gradient is shown in Supplementary Table 7. A coupled DAD-3000RS diode array detector (Dionex) collected UV absorption spectra in the range of 190-560 nm, a FLD-311 fluorescence detector (Dionex) collected fluorescence data at 275 nm excitation and 350 nm emission, and a MSQPlus mass spectrometer (Dionex) collected ESI mass spectra in positive and negative ion modes in the range of 100-1000 m/z. Total ICN, 4OH-ICN and camalexin amounts were quantified using standard curves of standards prepared in *cyp79B2 cyp79B3* seedling extract and integrated areas in the UV chromatographs at 260-nm for 4M-I (retention time [RT] = 7.7 min); 340-nm for ICN (RT = 11.5 min); 280-nm for ICN degradation product ICA-ME (RT = 9.5 min); and co-eluting 4OH-ICN degradation products 4OH-ICA and 4OH-ICA-ME (RT = 10.1 min); and 320 nm for camalexin (RT = 12.1 min). For Figure 1b, total camalexin amounts were quantified using integrated areas in the FLD chromatograph. For some experiments, 2.5 uL 200 μM indole butrytic acid (IBA; RT = 10.1 min) was added per mg sample dry weight instead of 4M-I. Relative amounts of ICN, 4OH-ICN, and amounts were quantified by dividing the peak areas at m/z 169 [M-H]-(ICN), 174 [M-H]-(ICA-ME), 176 [M-H]-(4OH-ICA), 190 [M-H]-(4OH-ICME), and 201 [M+H]+ (camalexin), by that of IBA (m/z 202 [M-H]-).

### Glucosinolate extraction and LC-DAD-FLD-MS

Glucosinolates were analyzed as desulfoglucosinolates as previously described by Kliebenstein *et al*. (2001) with some modifications. Briefly, a 96-well 0.45 μm PVDF filter plate (EMD Millipore, Billerica, MA) was charged with 45 mg DEAE Sephadex A25 (GE Heathcare) and 300 μL of water per well and equilibrated at room temp for 2 h. Prior to sample homogenization, the plate was centrifuged at 400xg for 1 min to remove the water. The homogenate was extracted with 500 μL 70% (v/v) aqueous methanol at 67.5°C for 10 min and centrifuged at 16,000xg for 2 min. Added to the supernatant was 3 μL of IS (1.25 mM sinigrin (Sigma-Aldrich) in 80% (v/v) ethanol) per mg sample dry weight. Extract was applied to and incubated on the ion exchanger for 10 min. The sephadex resin was washed three times with 70% (v/v) methanol, three times with distilled deionized water (ddH_2_O), and two times with 20 mM sodium acetate (pH 5). 20μL of 25 mg/mL aryl sulfatase (Type H1 from *Helix pomatia*, Sigma-Aldrich) was applied to and incubated on the sephadex resin at RT overnight (Hogge *et al*., 1988). The plate was centrifuged at 400xg for 1 min, and desulfoglucosinolates were eluted from the sephadex resin by two 100-μL washes with 60% (v/v) methanol and two 100-μL washes with ddH_2_O. Eluate volume was reduced to 250-350 μL using an evaporator. Samples were separated using the gradient shown in Supplementary Table 7. A coupled DAD-3000RS diode array detector, FLD-311 fluorescence detector (Dionex), and MSQPlus mass spectrometer collected UV

absorption spectra at 229-nm, fluorescence spectra at 275/350-nm (ex/em), and ESI mass spectra in positive/negative ion modes at 100-1000 m/z, respectively. Glucosinolates were quantified using integrated areas of desulfoglucosinolates in the UV chromatographs at 229-nm and published response factors (Clarke, 2010).

### Chromatin immunoprecipitation and (q)PCR

ChIP was performed as previously described by Chezem et al. (2017) with some modifications. Approximately two-hundred-and-ten 9-day-old seedlings were inoculated with *Pto* DC3000 *avrRpm1* to OD_600_ of 0.013 and co-treated with mock solution of DMSO (M) or 20 µM dexamethasone (D) for 9 hr. Following nuclear extraction, samples were sonicated in a Covaris S2 sonicator (Covaris, Woburn, MA) using 10% duty, 7% intensity, 200 cycles per burst for a total time of 11 min. Chromatin immunoprecipitation was performed using Anti-FLAG M2 Affinity Gel (Sigma-Aldrich). Beads were pre-treated with 0.1% (w/v) non-fat milk in 1X PBS and 0.5 mg/mL sheared salmon sperm DNA (Invitrogen). Following de-crosslinking, DNA samples were phenol-chloroform-extracted, diluted to the same OD_260_ concentration, and 1.5 µL was used in 15 µL PCR reaction. PCR analysis was performed on nuclear extracts prior to antibody incubation (input) and after ChIP. PCR conditions were as follows: 95°C for 3 min; 40 cycles of 95°C for 15 sec, 53°C for 15 sec, and 72°C for 1 min; 72°C for 5 min. Densitometric determination of signal intensity in each ChIP and input sample was calculated using ImageJ. Fold enrichment was determined by calculating the ratio of PCR product intensities in ChIP D/M to Input D/M. In cases where amplicons were absent, an arbitrary value of 10 was assigned. For *EPL2*, qPCR analysis was additionally performed to confirm absence of amplicons in ChIP samples. RLU counts at the 25^th^ cycle were used for quantification. Primer sequences are listed in Supplementary Table 6.

### Comparative genomics

All phylogenetic species trees were adapted from Koch and Kiefer (2005) and Couvreur et al. (2009). To generate novel phylogenetic maximum likelihood (ML) trees, sequences were aligned using MUSCLE in MEGA7 (Kumar *et al*., 2016) and JTT model (for CYP82C and LINE alignments) or Tamura-Nei model (for the *EPCOT3* alignment). Sequences for all genes with the description “non-LTR retrotransposon family (LINE)” (N=263) were batch-downloaded from TAIR (https://arabidopsis.org). Of these, sequences containing intact reverse transcriptase domains (“PGPDG”, “LIPK”, “FRPISL”, or “FADD” sequences; N=126) were used for subsequent phylogenetic analysis. Gaps were removed from the CYP82C alignment, leaving a total of 480 codons. *EPCOT3* alignments were visualized in JalView (http://www.jalview.org/; Waterhouse *et al*., 2009). Information on genomes used for synteny analysis is shown in Supplementary Table 8.

Selection estimates based on nonsynonymous-to-synonymous substitution ratios were calculated from the CYP82C ML tree (Supplementary Text 1). A Newick tree file was generated from this ML tree (Supplementary Figure 4b; Supplementary Table 2) and for Branch site models, branches were pre-defined. CodeML analysis in PAML (Yang, 2007) was then conducted with the following modified parameters: ncatG = 8; CodonFreq = 3. The M0 test was performed with model = 0 and NSsites = 0. The M1a null test was performed with model = 0 and NSsites = 1. A more stringent null test (fixed omega) was performed for each Branch site model to be tested (model = 2 and NSsites = 2), where omega was fixed to 1. Branch site models were then tested with unfixed omega. Likelihood ratio tests were performed by comparing critical values and degrees of freedom between each unfixed Branch site test and either the M1a test or the corresponding fixed-omega test. Pre-defined branches with *P* values less than 0.05 for both tests were regarded as under positive selection (Supplementary Figure 2).

The protein structure of CYP82C2 was generated using Intensive modeling mode in Phyre2 (http://www.sbg.bio.ic.ac.uk/phyre2/html/page.cgi?id=index; Kelley *et al*., 2015) and visualized in MacPyMOL (Schrödinger, LLC). Amino acid conservation was scored using the Bayesian Best model in Consurf (http://consurf.tau.ac.il/2016/; Ashkenazy *et al*., 2016).

### Bioinformatics

Coexpression data was obtained from ATTED-ii (http://atted.jp/; Obayashi et al., 2018). Mutual ranks less than 200 are indicative of strong co-expression (Obayashi et al., 2018). Epigenetics data was obtained from Roudier et al. (2011) and confirmed using data from Liu et al. (2018). Percent identity matrices were constructed from Clustal Omega Multiple Sequence Alignments (https://www.ebi.ac.uk/Tools/msa/clustalo/). Promoter alignment plots were generated using mVISTA (http://genome.lbl.gov/vista/mvista/submit.shtml; Frazer et al., 2004)

## Data and code availability

The authors declare that all data supporting the findings of this study are available within the manuscript and the Supplementary Information or are available from the corresponding authors upon request. Custom code to visualize homology models in MacPyMOL and conduct statistics in R are also available upon request.

## Acknowledgements

We thank J.L. Celenza for the *cyp79B2 cyp79B3* mutant. We thank E.S. Sattely for ICN/ICN-ME, 4OH-ICA/4OH-ICA-ME and camalexin standards. This work was supported by T32-GM007499 (to B.B.) and Elsevier/Phytochemistry Young Investigator Award (to N.K.C.).

## Author Contributions

B.B. and N.K.C. performed pathogen assays and ChIP-PCR experiments. N.K.C. performed RT-PCR experiments. B.B. and Y.K. profiled accessions and species. B.B. performed all other experiments. B.B. and N.K.C. interpreted the results and wrote the paper.

## Materials & Correspondence

Correspondence and material requests can be addressed to Brenden Barco.

