## Supplementary Texts, Figures, Tables for "Expansion of a core regulon by transposable elements promotes Arabidopsis chemical diversity and pathogen defense"

Affiliations:

(((((((Ta22L1:0.13595018,Ta22L2:0.11294348)0.5400:0.05991792,Ta22L3:0.12321666)0.1800:0.05249874,Ta22L4:0.14202430)0.2000:0.00628800,(Ta22:0.13407885,Ta22L5:0.20261792)0.2600:0.04059211)0.1600:0.08786300,(Ta22L6:0.16667474,(Ta22L7:0.00739203,Ta22L8:0.00708679)1.0000:0.09029017)0.4600:0.03821092)0.4000:0.06981106,EPL2:0.19695608)0.3800:0.03607816,(EPCOT3:0.03788569,EPL1:0.02923475)0.8200:0.06306515,Ta20:0.70226168);

**Supplementary Text 1.** Full phylogenetic maximum likelihood tree of *EPCOT3* and related TE sequences in Newick format. *A. thaliana* LINE *Ta20* was used as the outgroup.

((((((((((((('rf_3_AT2G25550.1_chr2:10874913..10880212_FORWARD_LENGTH_5300':0.00000000,'rf_3_AT2G31520.1_chr2:13421940..13427089_REVERSE_LENGTH_5150':0.00000000)0.7200:0.00000000,'rf_2_AT5G37665.1_chr5:14962141..14967533_REVERSE_LENGTH_5393':0.00671111)1.0000:0.04571846,'rf_2_AT1G35960.1_chr1:13390829..13396127_REVERSE_LENGTH_5299':0.08266884)0.3400:0.01128056,('rf_3_AT5G35076.1_chr5:13348106..13353283_FORWARD_LENGTH_5178':0.04332076,('rf_1_AT3G45550.1_chr3:16709445..16712000_REVERSE_LENGTH_2556':0.00258660,'rf_3_AT5G53775.1_chr5:21831518..21836800_REVERSE_LENGTH_5283':0.00659435)0.9000:0.01872966)0.5500:0.01579459)0.7900:0.03055492,('rf_3_AT1G41850.1_chr1:15626402..15630507_FORWARD_LENGTH_4106':0.07743050,('rf_2_AT1G30030.1_chr1:10531831..10534842_FORWARD_LENGTH_3012':0.01348099,('rf_3_AT2G14990.1_chr2:6475058..6480183_FORWARD_LENGTH_5126':0.01344875,'rf_3_AT4G10830.

1_chr4:6650724..6654712_FORWARD_LENGTH_3989':0.00000000)0.5100:0.00479300)0.9600:0.02441992)0.9300:0.08034886)1.0000:0.20403922,('rf_1_AT5G27845.1_chr5:9868348..9871836_FORWARD_LENGTH_3489':0.16894611,('rf_1_AT5G35413.1_chr5:13639182..13643273_REVERSE_LENGTH_4092':0.09544888,'rf_2_AT3G32043.1_chr3:13053130..13058318_REVERSE_LENGTH_5189':0.20548606)0.4000:0.01955888)0.9700:0.12862360)0.3500:0.03741481,(((('rf_3_AT2G06540.1_chr2:2592442..2597780_REVERSE_LENGTH_5339':0.08713438,'rf_3_Ta15_AT4G06497.1_chr4:3138929..3144238_REVERSE_LENGTH_5310':0.07019209)0.5600:0.03291158,'rf_2_AT2G12195.1_chr2:4894628..4898737_REVERSE_LENGTH_4110':0.05808295)0.8000:0.06621267,'rf_3_AT3G44705.1_chr3:16244952..16250205_FORWARD_LENGTH_5254':0.11624708)1.0000:0.10830430,('rf_3_1213_AT5G35331.1_chr5:13526418..13531682_REVERSE_LENGTH_5265':0.21262331,(('rf_1_AT2G12650.1_chr2:5172044..5177532_FORWARD_LENGTH_5489':0.13883339,'rf_2_AT4G04405.1_chr4:2171191..2174337_REVERSE_LENGTH_3147':0.11399091)0.9500:0.08000369,('rf_1_AT3G29205.1_chr3:11170646..11175547_REVERSE_LENGTH_4902':0.20317952,('rf_3_AT2G07160.1_chr2:2969531..2974603_FORWARD_LENGTH_5073':0.10200579,'rf_3_AT3G31420.1_chr3:12787914..12792017_REVERSE_LENGTH_4104':0.17777558)0.6600:0.05719808)0.7700:0.03846058)0.7900:0.07197741)0.5900:0.04854001)0.8800:0.06621759)'TA12/13/15_clade'0.5200:0.03866977,(TA16_AT5G41835:0.16355448,TA23_AT2G01840:0.16195931)1.0000:0.27216131)0.2300:0.03783439,((('rf_1_AT1G35390.1_chr1:13008911..13014607_FORWARD_LENGTH_5697':0.02045919,'rf_3_AT5G36005.1_chr5:14144014..14149194_FORWARD_LENGTH_5181':0.02969527)1.0000:0.21929416,'rf_1_AT3G43

573.1_chr3:15487248..15491948_FORWARD_LENGTH_4701':0.38166553)1.0000:0.18720906,(('rf_3_AT3G43622.1_chr3:15527180..15530149_REVERSE_LENGTH_2970':0.10674777,'rf_3_AT4G09710.1_chr4:6128688..6132598_FORWARD_LENGTH_3911':0.04818830)1.0000:0.16645767,('rf_1_AT2G05200.1_chr2:1883052..1887007_FORWARD_LENGTH_3956':0.26236977,(('rf_2_AT2G06290.1_chr2:2475752..2484523_FORWARD_LENGTH_8772':0.19669864,'rf_3_AT5G35535.1_chr5:13713460..13719312_FORWARD_LENGTH_5853':0.28402216)0.3000:0.04263875,(('rf_2_TA21_AT2G10820.1_chr2:4256189..4261900_FORWARD_LENGTH_5712':0.20317936,'rf_3_AT4G06644.1_chr4:3795731..3801568_FORWARD_LENGTH_5838':0.27936915)0.5300:0.07234094,((('rf_1_AT4G04000.1_chr4:1920423..1925005_FORWARD_LENGTH_4583':0.15778149,'rf_3_TA18_AT5G24915.1_chr5:8571956..8577856_FORWARD_LENGTH_5901':0.04492595)0.9400:0.08067966,'rf_1_AT5G28545.1_chr5:10547746..10553612_REVERSE_LENGTH_5867':0.14195996)0.4300:0.05521487,(('rf_1_AT3G43575.1_chr3:15493460..15497791_REVERSE_LENGTH_4332':0.09996502,'rf_1_AT4G01490.1_chr4:631136..636899_FORWARD_LENGTH_5764':0.02817618)1.0000:0.11167005,('rf_3_TA20_AT5G28053.1_chr5:10050551..10056352_FORWARD_LENGTH_5802':0.09891750,('rf_2_AT5G43415.1_chr5:17440088..17445889_FORWARD_LENGTH_5802':0.04381224,'rf_3_AT2G11240.1_chr2:4477686..4483748_REVERSE_LENGTH_6063':0.15601927)0.8700:0.05758397)0.6500:0.06299815)0.5600:0.03132201)0.3100:0.04616756)0.0700:0.01771117)0.7200:0.08251906)0.1700:0.03848175)1.0000:0.21594302)'TA18/20_clade'0.3300:0.07588287)0.2400:0.08839073,((Ta22_AT5G28523:0.23366400,Ta22L_AT2G45230:0.08479209)1.0000:0.31170890,((((('rf_1_AT2G16560.1_chr2:7173003..71

76689_FORWARD_LENGTH_3687':0.04344013,'rf_3_AT2G11820.1_chr2:4752782..4756899_REVERSE_LENGTH_4118':0.05374966)0.2300:0.00686565,'rf_2_AT1G45140.1_chr1:17071358..17074556_REVERSE_LENGTH_3199':0.06784932)0.2700:0.02222684,('rf_1_AT2G12640.1_chr2:5154645..5158744_REVERSE_LENGTH_4100':0.07003851,'rf_3_AT1G43270.1_chr1:16319951..16324362_FORWARD_LENGTH_4412':0.03348406)0.7900:0.01262848)0.2800:0.03655787,'rf_1_AT1G47910.1_chr1:17656860..17660288_FORWARD_LENGTH_3429':0.00000000)1.0000:0.22740430,(('rf_2_AT3G47875.1_chr3:17663475..17666891_REVERSE_LENGTH_3417':0.05941429,'rf_3_AT5G35495.1_chr5:13694865..13698386_FORWARD_LENGTH_3522':0.13222152)1.0000:0.13757993,('rf_3_AT2G15540.1_chr2:6778957..6783018_REVERSE_LENGTH_4062':0.02842552,('rf_3_TA25_AT2G05780.1_chr2:2193600..2196125_REVERSE_LENGTH_2526':0.04838410,('rf_1_AT2G12670.1_chr2:5182756..5185602_FORWARD_LENGTH_2847':0.05856511,'rf_1_AT2G16420.1_chr2:7116052..7119987_FORWARD_LENGTH_3936':0.04899137)0.7600:0.02245468)0.6200:0.02891924)1.0000:0.10660815)0.7100:0.08056846)'TA25_clade'0.9400:0.11381873)0.5000:0.08280714)0.2200:0.07510435,(((((('rf_1_TA11/ATLN54_AT3G24675.1_chr3:9011349..9015500_REVERSE_LENGTH_4152':0.11172102,'rf_1_AT5G28405.1_chr5:10350541..10354653_REVERSE_LENGTH_4113':0.11175482)0.9700:0.10342573,'rf_3_AT3G28945.1_chr3:10970273..10975947_REVERSE_LENGTH_5675':0.11864968)0.8800:0.10670303,('rf_1_TA24_AT1G24640.1_chr1:8729280..8732681_FORWARD_LENGTH_3402':0.11845491,'rf_1_AT1G35146.1_chr1:12856198..12860244_REVERSE_LENGTH_4047':0.18796192)0.8400:0.09439293)0.8100:0.08328853,'rf_1_AT5G04235.1_chr5:1164287..1168411_REVERSE_LEN

GTH_4125':0.32079664)0.5600:0.08927846,(('rf_1_AT2G13490.1_chr2:5623788..5627
656_FORWARD_LENGTH_3869':0.25470559,'rf_1_AT2G17910.1_chr2:7778495..7782
529_FORWARD_LENGTH_4035':0.03569312)0.8800:0.04224443,('rf_2_AT2G13460.1
_chr2:5600306..5605593_REVERSE_LENGTH_5288':0.11939177,(('rf_1_AT2G15250.
1_chr2:6619688..6624973_REVERSE_LENGTH_5286':0.14812527,'rf_3_AT3G33565.
1_chr3:14060883..14064624_FORWARD_LENGTH_3742':0.27465770)0.2600:0.02651
071,('rf_1_AT3G57586.1_chr3:21322564..21329357_FORWARD_LENGTH_6794':0.12
396726,('rf_1_AT2G41580.1_chr2:17339804..17343088_FORWARD_LENGTH_3285':
0.10563151,'rf_2_AT5G18633.1_chr5:6205813..6210644_FORWARD_LENGTH_4832':
0.07476436)0.7700:0.04478752)0.2900:0.01978951)0.5900:0.05073149)0.5700:0.0441
0362)0.9800:0.17570359)'TA11/24_clade'0.7900:0.08643175)0.9300:0.17301557,(LOR
F2_O00370:1.05556441,('rf_3_AT3G43175.1_chr3:15173888..15177508_REVERSE_L
ENGTH_3621':0.10267667,((('rf_1_AT1G58020.1_chr1:21450213..21455851_REVERS
E_LENGTH_5639':0.14544094,'rf_2_AT5G43105.1_chr5:17304711..17308952_FORW
ARD_LENGTH_4242':0.41083239)0.1600:0.03514594,('rf_1_AT3G32110.1_chr3:1309
7952..13103951_FORWARD_LENGTH_6000':0.09101707,'rf_1_AT3G45253.1_chr3:16
589291..16594622_REVERSE_LENGTH_5332':0.10438749)0.6000:0.03429819)0.100
0:0.04018045,('rf_1_AT2G07650.1_chr2:3222935..3225703_REVERSE_LENGTH_276
9':0.13153439,('rf_3_AT4G08830.1_chr4:5623753..5627210_FORWARD_LENGTH_34
58':0.18065264,(('rf_1_AT5G36935.1_chr5:14574600..14584700_FORWARD_LENGT
H_10101':0.19330207,'rf_3_TA28_AT5G07505.1_chr5:2374053..2377382_FORWARD
_LENGTH_3330':0.11307642)0.2600:0.04134002,('rf_1_AT4G15590.1_chr4:8900245..

8907255_REVERSE_LENGTH_7011':0.08499607,('rf_2_AT2G31080.1_chr2:13223059..13227048_REVERSE_LENGTH_3990':0.08293752,'rf_3_AT1G43250.1_chr1:16311524..16316838_FORWARD_LENGTH_5315':0.10773575)0.2800:0.02143400)0.5200:0.04329528)0.0200:0.03499223)0.0300:0.04232329)0.0400:0.02942516)0.1500:0.05161331)'TA28_clade'0.9700:0.27709268)0.5400:0.17124347,((((('rf_1_AT4G26360.1_chr4:13328763..13332375_REVERSE_LENGTH_3613':0.00000000,'rf_2_AT5G40605.1_chr5:16259229..16264465_FORWARD_LENGTH_5237':0.01370783)1.0000:0.08004083,'rf_3_AT1G25430.1_chr1:8927732..8933058_REVERSE_LENGTH_5327':0.11536533)0.6100:0.05527848,('rf_1_AT2G15720.1_chr2:6845385..6848332_REVERSE_LENGTH_2948':0.13826988,'rf_1_AT5G55896.1_chr5:22628017..22633342_REVERSE_LENGTH_5326':0.05665341)0.9400:0.08538116)0.5800:0.04737030,('rf_1_AT1G31100.1_chr1:11097034..11100418_REVERSE_LENGTH_3385':0.20708908,('rf_3_AT4G03920.1_chr4:1855717..1861200_FORWARD_LENGTH_5484':0.19553908,('rf_1_AT3G26614.1_chr3:9784890..9787571_REVERSE_LENGTH_2682':0.04548152,'rf_1_AT5G47815.1_chr5:19358618..19363928_FORWARD_LENGTH_5311':0.11537425)0.7200:0.05256428)0.2700:0.02457562)0.1300:0.03347328)0.9900:0.16186004,((((('rf_1_AT4G02490.1_chr4:1097945..1100011_REVERSE_LENGTH_2067':0.02109420,'rf_1_AT5G13475.1_chr5:4320277..4326003_FORWARD_LENGTH_5727':0.00000000)0.9700:0.07264702,'rf_2_AT1G10160.1_chr1:3328971..3333470_FORWARD_LENGTH_4500':0.12144899)0.7500:0.04683720,'rf_1_AT4G06523.1_chr4:3302313..3305957_FORWARD_LENGTH_3645':0.20128085)0.7800:0.08567355,'rf_2_AT5G07215.1_chr5:2257475..2262962_REVERSE_LENGTH_5488':0.24052648)1.0000:0.25198056,('rf_3_AT5G27905.1_chr5:

9925219..9930411_FORWARD_LENGTH_5193':0.41913841,('rf_3_AT5G27900.1_chr5:9913203..9918023_FORWARD_LENGTH_4821':0.86672166,((((('rf_1_AT3G05415.1_chr3:1555537..1560588_REVERSE_LENGTH_5052':0.02446292,'rf_2_AT2G05980.1_chr2:2313803..2317933_REVERSE_LENGTH_4131':0.06863152)1.0000:0.20664589,'rf_3_AT2G18820.1_chr2:8150995..8156099_FORWARD_LENGTH_5105':0.11506820)0.3200:0.04136140,('rf_2_AT2G06560.1_chr2:2611034..2616254_FORWARD_LENGTH_5221':0.12556210,'rf_2_AT3G43357.1_chr3:15307272..15312506_FORWARD_LENGTH_5235':0.10224139)0.9600:0.14647336)0.2800:0.05486522,('rf_3_AT2G28980.1_chr2:12449336..12454356_REVERSE_LENGTH_5021':0.05000389,('rf_3_AT1G47860.1_chr1:17620677..17630270_REVERSE_LENGTH_9594':0.01288227,'rf_3_AT2G14430.1_chr2:6134010..6139121_FORWARD_LENGTH_5112':0.01414129)1.0000:0.06301752)1.0000:0.11827546)0.0600:0.01918239,((('rf_2_AT3G29778.1_chr3:11660602..11663681_FORWARD_LENGTH_3080':0.09487799,'rf_3_AT4G06630.1_chr4:3754366..3757659_FORWARD_LENGTH_3294':0.02846571)0.7300:0.06116856,'rf_2_AT3G62725.1_chr3:23203792..23207565_REVERSE_LENGTH_3774':0.04686265)0.9800:0.12628687,('rf_1_AT3G44650.1_chr3:16209849..16212164_REVERSE_LENGTH_2316':0.11418628,(((('rf_2_AT1G32890.1_chr1:11914967..11919845_REVERSE_LENGTH_4879':0.04941054,'rf_3_AT5G35725.1_chr5:13885445..13890247_FORWARD_LENGTH_4803':0.07925394)0.9600:0.06046151,('rf_2_AT2G05550.1_chr2:2036684..2040701_REVERSE_LENGTH_4018':0.33269760,'rf_3_AT3G44425.1_chr3:16057731..16060385_FORWARD_LENGTH_2655':0.04428116)0.0300:0.00000000)0.1200:0.02079835,'rf_1_AT1G31030.1_chr1:11064583..11067201_REVERSE_LENGTH_2619':0.07323065)0.2

000:0.01677079,('rf_3_AT3G43315.1_chr3:15271837..15275043_REVERSE_LENGTH_3207':0.17810344,('rf_3_AT5G39245.1_chr5:15718364..15722158_REVERSE_LENGTH_3795':0.08830726,((('rf_1_AT5G39862.1_chr5:15961511..15964681_FORWARD_LENGTH_3171':0.00000000,'rf_3_AT2G01550.1_chr2:243918..249049_REVERSE_LENGTH_5132':0.00680847)1.0000:0.08119811,'rf_3_AT3G25485.1_chr3:9236887..9240548_REVERSE_LENGTH_3662':0.12456183)0.3600:0.03210122,('rf_3_AT2G19100.1_chr2:8269781..8274895_REVERSE_LENGTH_5115':0.09857376,('rf_1_AT5G26582.1_chr5:9388756..9392082_REVERSE_LENGTH_3327':0.04743447,('rf_1_AT3G14517.1_chr3:4869841..4872139_REVERSE_LENGTH_2299':0.10131591,'rf_2_AT2G23880.1_chr2:10165992..10170262_REVERSE_LENGTH_4271':0.07848497)0.7600:0.07164797)0.2800:0.02129676)0.2100:0.02896343)0.0100:0.01781385)0.0500:0.02079464)0.0400:0.02496510)0.5100:0.05069067)0.7300:0.08628325)0.1500:0.02649459)0.2900:0.06263123)0.8200:0.13240621)0.3800:0.11430363)0.3600:0.07065795)0.9600:0.29675307)
;

**Supplementary Text 2.** Full phylogenetic maximum likelihood tree of *A. thaliana* LINE reverse transcriptase proteins in Newick format.

**A**

| | | flg22 | | *Psta* | |
|---|---|---|---|---|---|
| | | fold change | *p*-value | fold change | *p*-value |
| | *CYP79B2* | 29.53 | 0.00E+00 | 875.50 | 2.49E-08 |
| | *CYP79B3* | 3.50 | 1.09E-10 | 74.73 | 2.85E-04 |
| 4OH-ICN | *CYP71A12* | 16.91 | 1.66E-10 | 76.06 | 4.16E-05 |
| | *FOX1* | 100.00 | 0.00E+00 | 19.14 | 2.00E-04 |
| | *CYP82C2* | 13.41 | 3.21E-06 | 642.21 | 8.90E-04 |
| camalexin | *CYP71A13* | 100.00 | 0.00E+00 | 590.44 | 2.03E-04 |
| | *CYP71B15* | 4.21 | 3.93E-14 | 591.30 | 7.59E-05 |
| 4M-I3M | *CYP83B1* | 10.18 | 4.35E-18 | 26.47 | 4.85E-04 |
| | *SUR1* | 13.76 | 7.32E-09 | 3.92 | 1.99E-04 |
| | *CYP81F2* | 100.00 | 0.00E+00 | 3.34 | 0.007 |

**D**

| | | 1 | 2 | 3 | 4 | 5 |
|---|---|---|---|---|---|---|
| CYP71A12 | 1 | | | | | |
| FOX1 | 2 | **1.4** | | | | |
| CYP82C2 | 3 | **7.5** | **22.1** | | | |
| CYP71A13 | 4 | **20.2** | **11.6** | **65.5** | | |
| CYP71B15 | 5 | **13.9** | **5.2** | **188.3** | **1.7** | |
| WRKY33 | 6 | 314.5 | **163.3** | 851.6 | 667.5 | 328.7 |

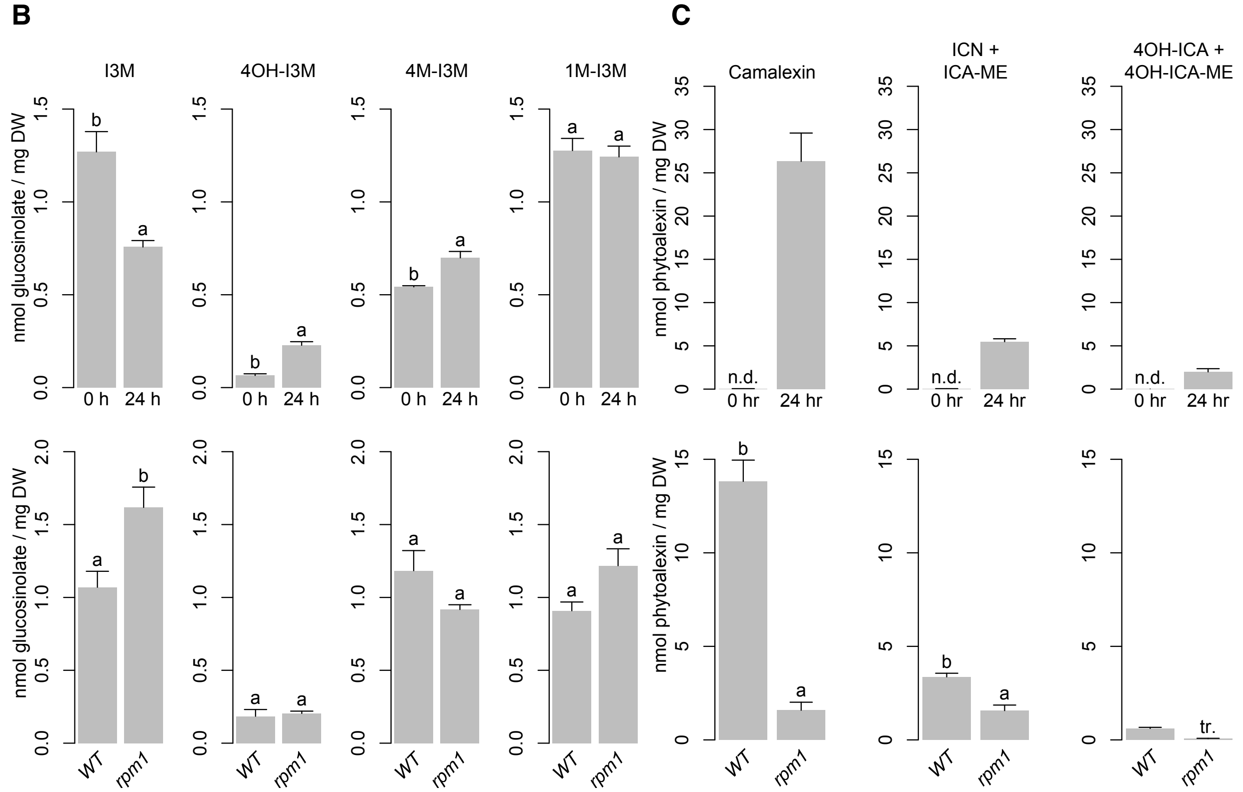

**Supplementary Figure 1. Defense responses in wild-type elicited plants.** (Supports Figure 1)

**a.** qPCR analysis of metabolite biosynthetic genes upregulated in response to 3-hr elicitation with 1 μM flg22 or 12-hr inoculation with the bacterial pathogen *Psta*. Flg22-

treated data obtained from Denoux *et al*. (2008). *Psta*-treated data represents the mean of 3 to 6 independent experiments, four biological replicates. ($P < 0.05$, two-tailed $t$ test).

**b-c.** HPLC-DAD analysis of I3M, 4OH-I3M, and 4M-I3M (**b**) and camalexin, ICN, and 4OH-ICN (**c**) in seedlings inoculated with *Psta* for 0 or 24 hr (top) and 24 hr (bottom). Data represent mean ± SE of four biological replicates. Different letters in denote statistically significant differences ($P < 0.05$, two-tailed $t$-test). DW, dry weight; n.d., not detected; tr., trace. Source data are provided as a Source Data file.

**d.** Co-expression (mutual rank) matrix between 4OH-ICN and camalexin biosynthetic genes. Mutual ranks less than 200 (in bold) are indicative of strong co-expression (Obayashi *et al*., 2018).

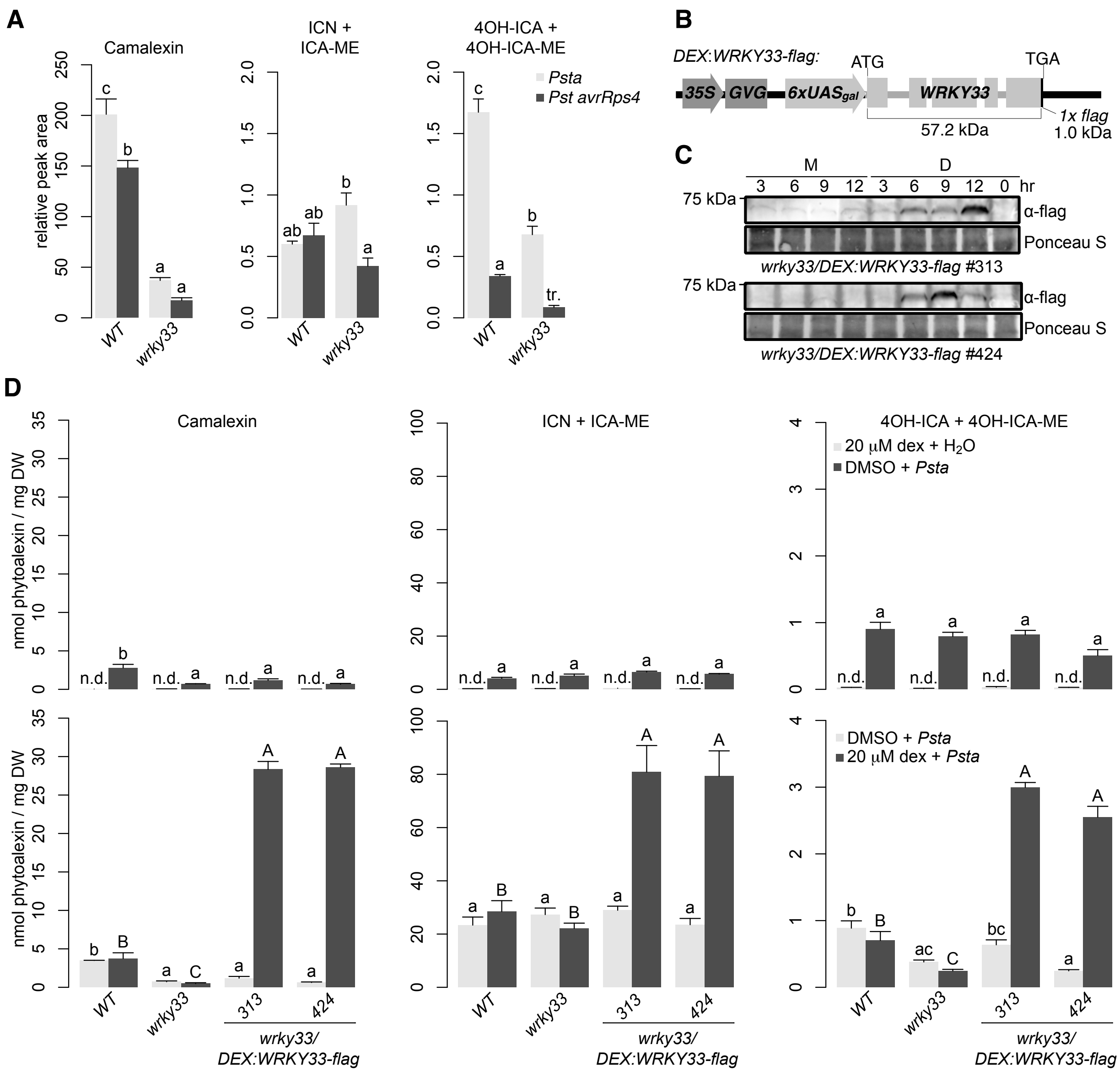

**Supplementary Figure 2.** (Supports Figure 2)

**a.** LC-DAD-MS of camalexin, ICN, and 4OH-ICN in seedlings inoculated with *Psta* or *Pst avrRps4* for 24 hr. Data represent mean ± SE of 3-4 replicates. Experiments were performed twice, producing similar results.

**b.** Schematic of the *DEX:WRKY33-flag* construct. Arrows indicate promoter elements; dark gray box, glucocorticoid-regulated transcription factor (GVG); light gray boxes,

*WRKY33* exons; and black box, 1x *flag* epitope. Only *WRKY33* and *flag* sequences are drawn to scale.

**c.** Immunoblot analysis of WRKY33-flag protein in seedlings co-treated with 20 µM dex (D) or mock (M, 0.5% DMSO) and *Psta* for 0, 3, 6, 9 and 12 hr.

**d.** HPLC-DAD analysis of camalexin, ICN and 4OH-ICN in seedlings following 24 hr of the indicated treatments. Data represent mean ± SE of 3-4 replicates.

Different letters in (a,d) denote statistically significant differences ($P < 0.05$, one-factor ANOVA coupled to Tukey's test). DW, dry weight; n.d., not detected; tr., trace.

For (**a,c-d**), source data are provided as a Source Data file.

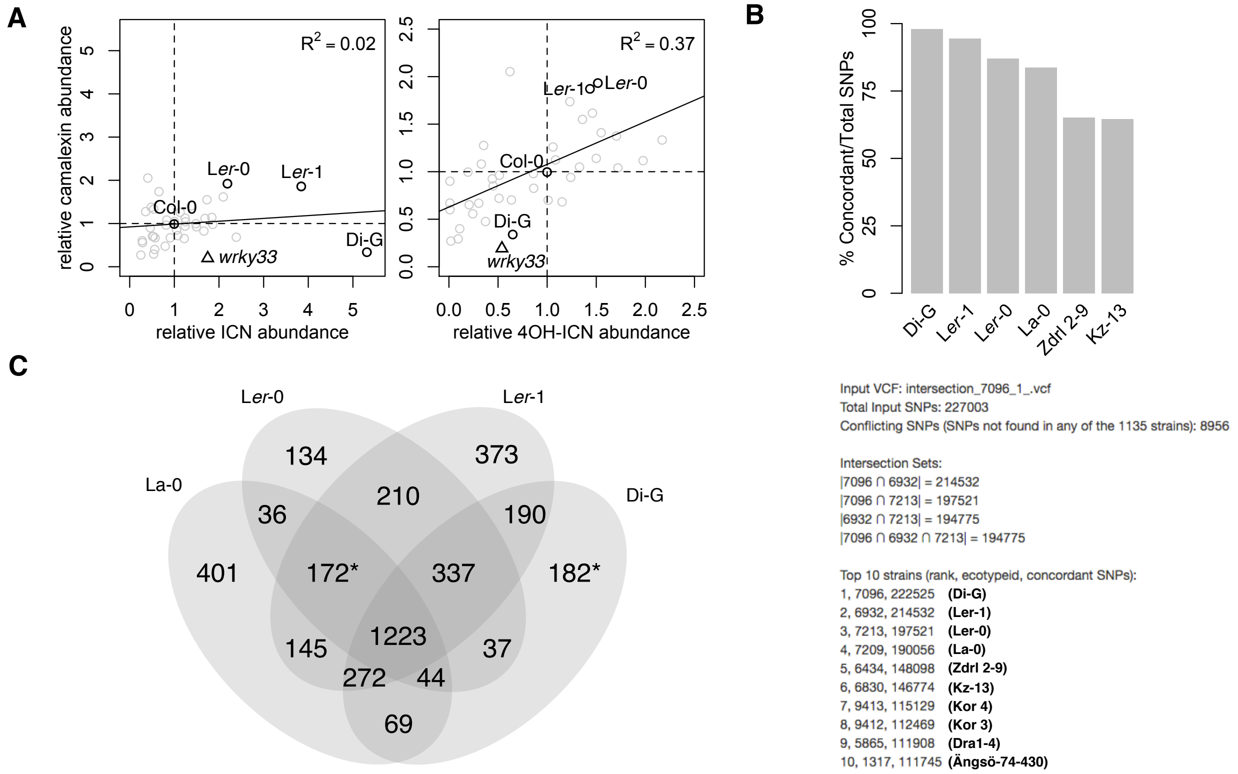

**Suppementary Figure 3.** (Supports Figure 2)

**a.** Scatterplot of camalexin and ICN (left) or camalexin and 4OH-ICN (right) levels in seedlings inoculated with *Psta* for 24 hr. Circles denote *A. thaliana* natural accessions (N = 40) and the triangle denotes the Col-0 *wrky33* mutant. Data represent mean of 2-4 replicates. Source data are provided as a Source Data file.

**b.** SNP details from the 1001 Genomes Strain ID tool using the published Di-G genome sequence variant file as input (1001 Genomes Consortium, 2016). Names of accessions added in bold.

**c.** Venn diagram of genes differentially mutated to high effect between Di-G, La-0, L*er*-0, and L*er*-1. Asterisks denotes genes used for subsequent GO term analysis**.**

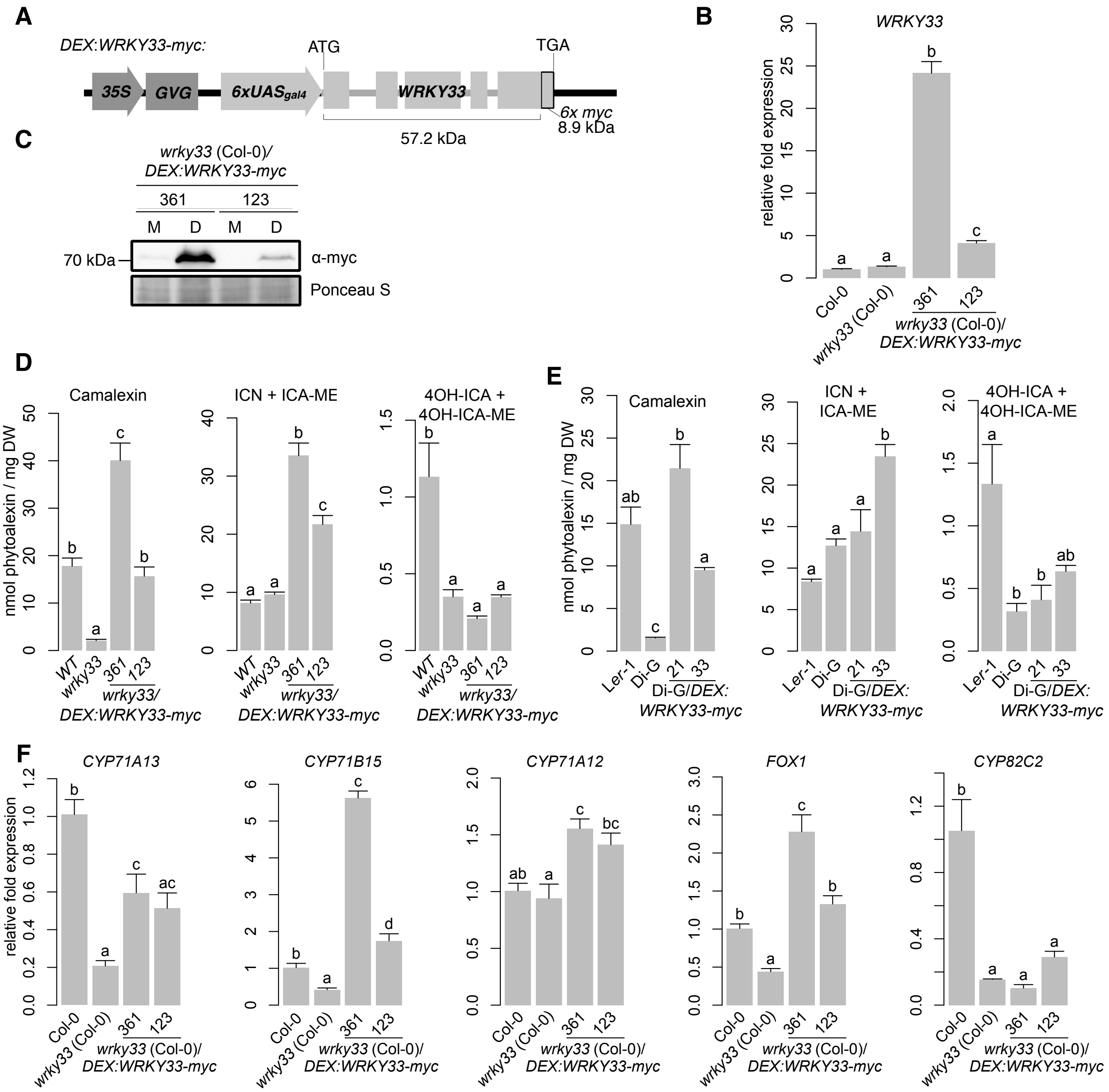

**Supplementary Figure 4.** (Supports Figure 2)

**a.** Schematic representation of the *DEX:WRKY33-myc* construct, consisting of the *WRKY33* gene fused to a 6x *myc* tag and driven by a two-component glucocorticoid-inducible promoter. Arrows indicate promoter elements, dark grey box indicates the glucocorticoid-regulated transcription factor (*GVG*), light gray boxes indicate the five

*WRKY33* exons, and the outlined box indicates the 6x myc epitope. Only *WRKY33* and *myc* sequences are drawn to scale.

**b.** qPCR analysis of *WRKY33* in seedlings inoculated with 20 µM dex and *Psta* for 12 hr.

**c.** Immunoblot analysis of WRKY33-flag protein in seedlings co-treated with 20 µM dex (D) or mock (M, 0.5% DMSO) and *Psta* for 6 hr.

**d-e.** HPLC-DAD analysis of camalexin, ICN and 4OH-ICN in seedlings co-treated with 20 µM dex and *Psta* for 24 hr. DW, dry weight.

**f.** qPCR analysis of *CYP71A13, CYP71B15, CYP71A12, FOX1,* and *CYP82C2* in seedlings inoculated with 20 µM dex and *Psta* for 12 hr.

Data in (**b**,**d-f**) represent mean ± SE of 3-4 replicates and different letters denote statistically significant differences ($P < 0.05$, one-factor ANOVA coupled to Tukey's test). Experiments in (b-f) were performed twice, producing similar results.

For (**b-f**), source data are provided as a Source Data file.

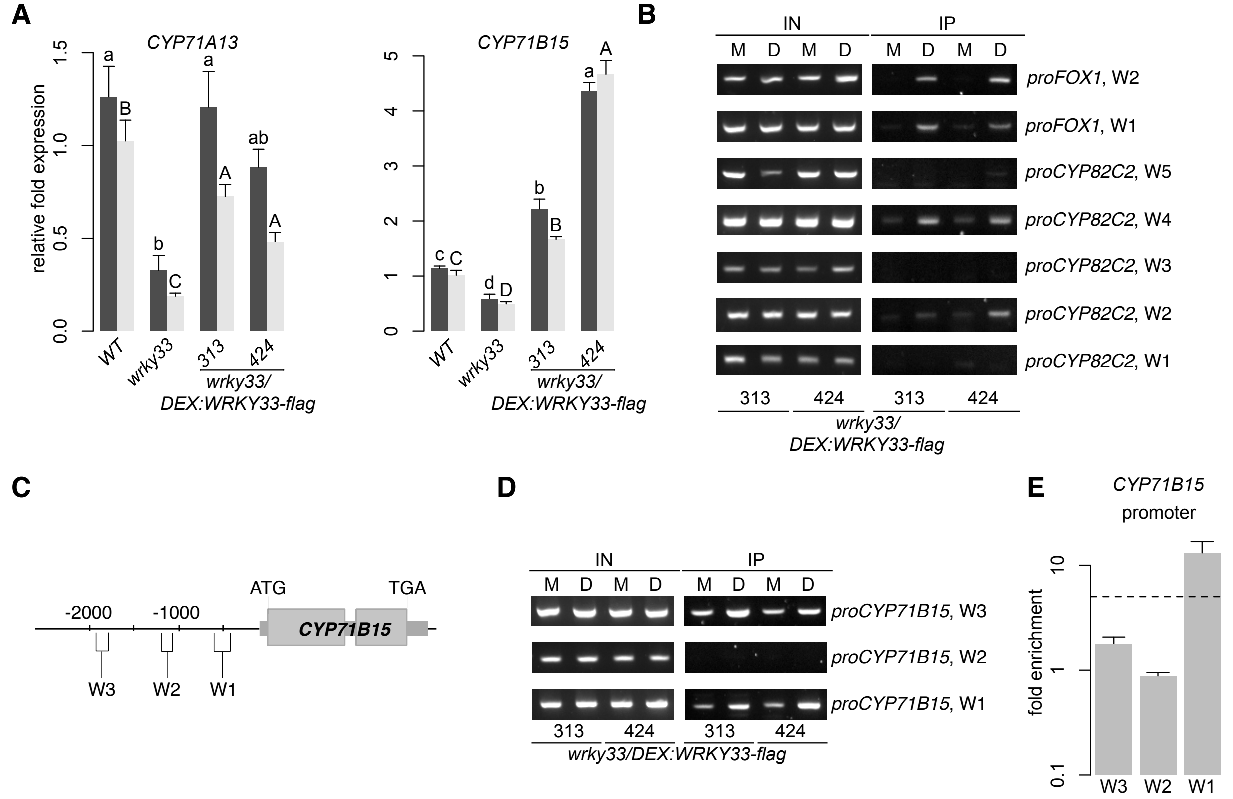

**Supplementary Figure 5. Functional characteristics of DEX:WRKY33-flag.**

(Supports Figure 3)

**a.** qPCR analysis of camalexin biosynthetic genes in seedlings inoculated with 20 µM dex and *Psta* for 9 and 12 hr. Different letters denote statistically significant differences ($P < 0.05$, one-factor ANOVA coupled to Tukey's test). Lowercase and uppercase letters denote comparisons across 9 and 12 hr timepoints, respectively. Data represent the mean ± SE of 4-6 replicates.

**b.** ChIP-PCR images of W-box-containing regions upstream of *FOX1* and *CYP82C2* in *wrky33/DEX:WRKY33-flag* plants co-treated with 20 µM dex (D) or mock solution (0.5% DMSO, M) and *Psta* for 9 hr.

**c.** Schematic of the *CYP71B15* locus, highlighting nt positions of W-box-containing regions.

**d.** ChIP-PCR images of W-box-containing regions in (**c**) in *wrky33/DEX:WRKY33-flag* plants co-treated with 20 µM dex (D) or mock solution (M, 0.5% DMSO) and *Psta* for 9 hr.

**e.** ChIP-PCR analysis of (**d**). Data represent mean ± SE of four replicates. Dashed line represents the 5-fold cutoff between weak and strong TF-DNA interactions.

For (**a-b**,**d-e**), source data are provided as a Source Data file.

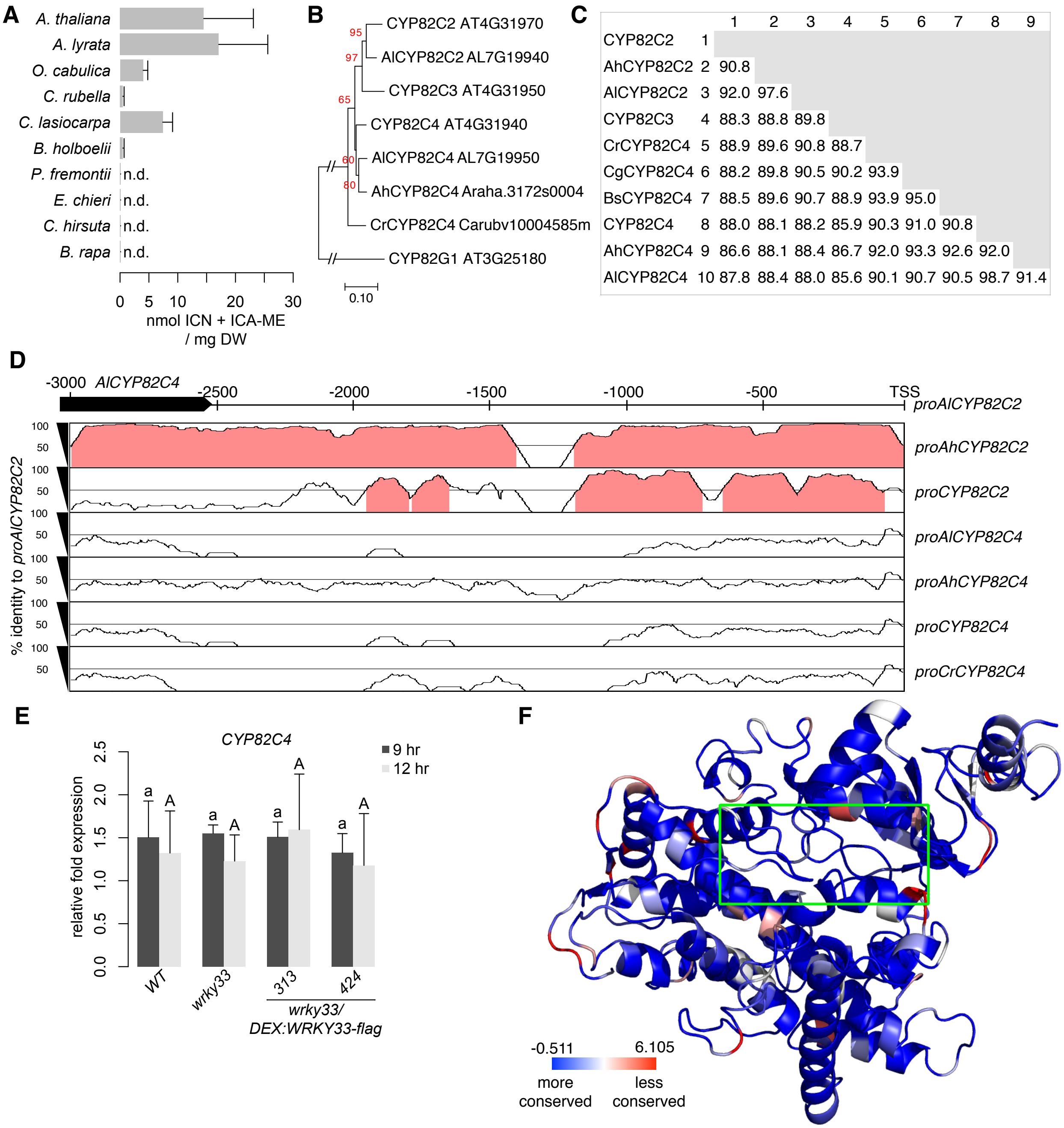

**Supplementary Figure 6.** (Supports Figure 4)

**a.** HPLC-DAD analysis of ICN in seedlings inoculated with *Psta* for 30 hr. Data represent mean ± SE of three independent experiments (n = 4 replicates), each with *A.*

*thaliana* as a positive control. Experiments were performed twice, producing similar results. ICA are breakdown products of 4OH-ICN. DW, dry weight; n.d., not detected.

**b.** Phylogenetic maximum likelihood tree of CYP82C family protein sequences in ICN-synthesizing species. Bootstrap values (N=100 replicate trees) are shown in red at the nodes. Scale bar represents 0.1 nucleotide substitutions per site. *AhCYP82C2* has a sequencing gap and was thus removed from analysis. *Al, Arabidopsis lyrata; Ah, Arabidopsis halleri*; *Cr, Capsella rubella*.

**c.** Percent identity matrix for encoded CYP82C enzymes in ICN-synthesizing species.

**d**. mVISTA plot of *AlCYP82C2* upstream sequence, indicating nt positions of conserved regions (≥70% sequence identity; pink) among homologous sequences. Also indicated is position of neighboring gene *AlCYP82C4* (black arrow). TSS, transcriptional start site; *Al, Arabidopsis lyrata; Ah, Arabidopsis halleri*; *Cr, Capsella rubella.*

**e.** qPCR analysis of *CYP82C4* in seedlings inoculated with 20 µM dex and *Psta* for 9 and 12 hr. Different letters denote statistically significant differences ($P < 0.05$, one-factor ANOVA coupled to Tukey's test). Lowercase and uppercase letters denote comparisons across 9 and 12 hr timepoints, respectively. Data represent the mean ± SE of 4-6 replicates.

**f.** Ribbon diagram of the homology model of CYP82C2. Amino acid residues are colored according to the conservation score at each residue. Boxed in green is the putative active site.

For (**a**,**e**), source data are provided as a Source Data file.

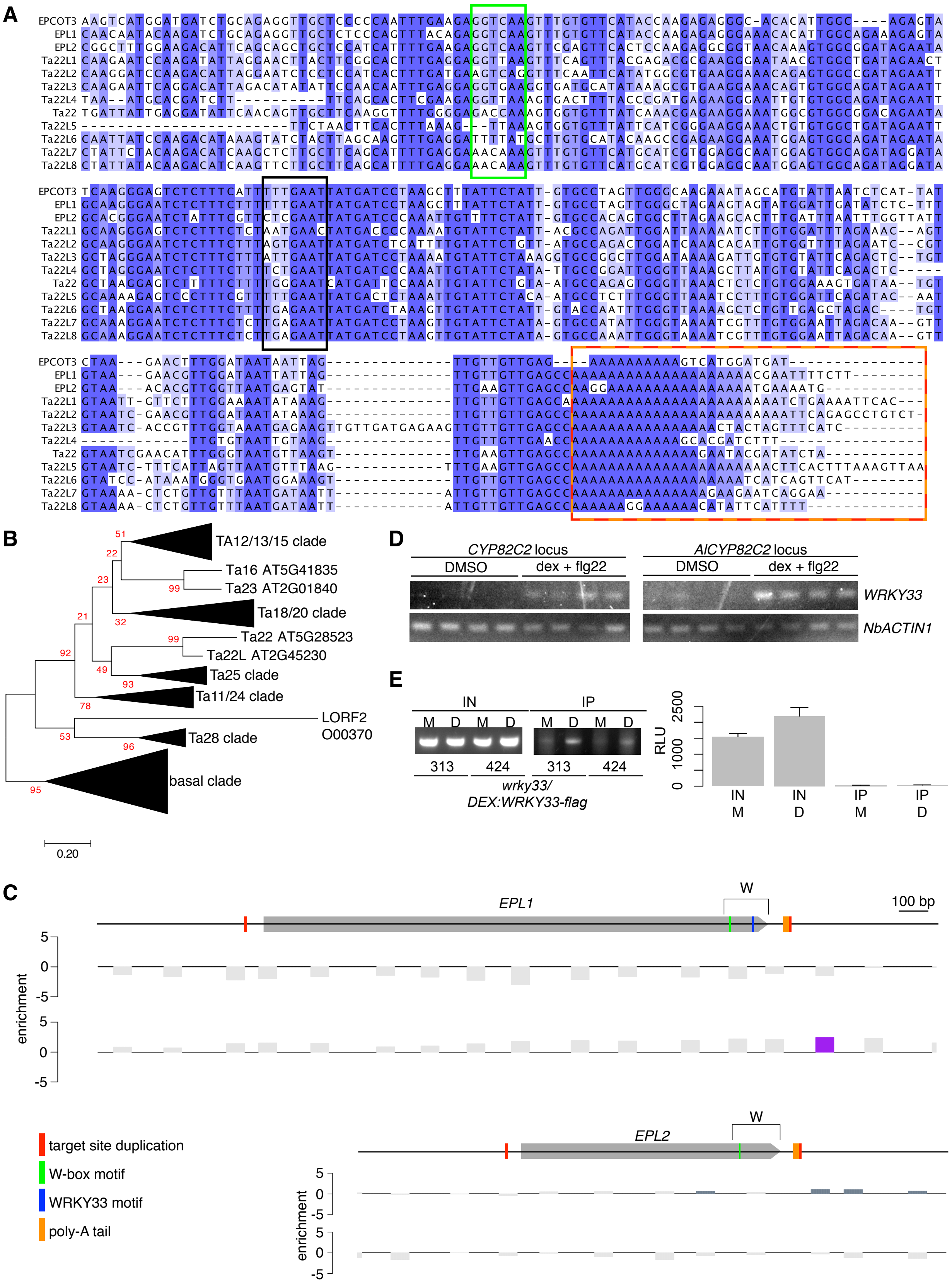
A
EPCOT3
EPL1
EPL2
Ta22L1
Ta22L2
Ta22L3
Ta22L4
Ta22
Ta22L5
Ta22L6
Ta22L7
Ta22L8
B
TA12/13/15 clade
Ta16 AT5G41835
Ta23 AT2G01840
Ta18/20 clade
Ta22 AT5G28523
Ta22L AT2G45230
Ta25 clade
Ta11/24 clade
LORF2
O00370
Ta28 clade
basal clade
0.20
D
CYP82C2 locus
AlCYP82C2 locus
DMSO
dex + flg22
WRKY33
NbACTIN1
E
IN
IP
M D M D
313
424
wrky33/
DEX:WRKY33-flag
RLU
IN M
IN D
IP M
IP D
C
EPL1
W
100 bp
enrichment
EPL2
target site duplication
W-box motif
WRKY33 motif
poly-A tail

**Supplementary Figure 7.** (Supports Figure 5)

**a.** Nucleotide alignment between *EPCOT3* and homologous sequences of related TEs. Also indicated are W-boxes (green box), WRKY33-specific motifs (black box), poly-A tails (red box), and 3' target site duplication (orange box).

**b.** Phylogenetic maximum likelihood tree of *A. thaliana* LINE reverse transcriptase proteins. *Homo sapiens* LINE1 ORF2 was used as the outgroup. Bootstrap values (N=100 replicate trees) are shown in red at the nodes. Scale bar represents 0.2 nucleotide substitutions per site. A more detailed tree is available as **Supplementary Text 2**.

**c.** Epigenetic map of *EPL1* (top) and *EPL2* (bottom), indicating nt positions of significant amounts of HK4me2 (blue-gray bars), and H3K27me3 (purple bars). Gray bars denote background amounts.

**d.** RT-PCR images of *WRKY33* and *NbACTIN1* expression in *N. benthamiana* leaves co-transfected with *DEX:WRKY33-flag* and the *CYP82C2* or *AlCYP82C2* locus and incubated for 15 hr with either mock solution (0.5% DMSO) or 20 µM dex plus 1 µM flg22.

**e.** (Top) ChIP-PCR image of W-box-containing regions within *EPL1* and (Bottom) ChIP-qPCR analysis of *EPL2* in *wrky33/DEX:WRKY33-flag* plants co-treated with 20 µM dex (D) or mock solution (0.5% DMSO, M) and *Psta* for 9 hr. Source data are provided as a Source Data file.

| description | TAIR ID | Landsberg state | Di-G state |
|---|---|---|---|
| Probable serine/threonine-protein kinase PBL28 | AT1G24030 | stop gained | WT |
| Disease resistance protein (TIR-NBS-LRR class) family | AT1G56520 | frameshift variant | WT |
| Disease resistance protein (TIR-NBS-LRR class) family | AT3G44630 | stop lost | WT |
| Myb/SANT-like DNA-binding domain protein | AT4G02550 | splice acceptor variant | WT |
| Putative defensin-like protein 28 | AT4G14272 | start lost | WT |
| Aquaporin NIP1-1 | AT4G19030 | frameshift variant | WT |
| Disease resistance protein (TIR-NBS-LRR class) | AT4G36140 | frameshift variant, start lost | WT |
| At5g03320 | AT5G03320 | splice donor variant | WT |
| Defensin-like protein 141 | AT5G47175 | frameshift variant | WT |
| Putative defensin-like protein 20 | AT5G52605 | frameshift variant | WT |
| Probable disease resistance protein | AT1G59620 | WT | frameshift variant |
| MLO-like protein 6 | AT1G61560 | WT | frameshift variant |
| PHLOEM PROTEIN 2-LIKE A5 | AT1G65390 | WT | frameshift variant, stop lost |
| Disease resistance protein (TIR-NBS-LRR class) family | AT1G65850 | WT | frameshift variant |
| (R)-mandelonitrile lyase-like | AT1G73050 | WT | frameshift variant |
| **Probable WRKY transcription factor 33** | **AT2G38470** | **WT** | **stop gained** |
| Endochitinase At2g43610 | AT2G43610 | WT | frameshift variant, start lost |
| Protein TIFY 6B | AT3G17860 | WT | frameshift variant |
| Disease resistance protein RPP13 | AT3G46530 | WT | frameshift variant |
| Ankyrin repeat family protein | AT4G03450 | WT | frameshift variant, stop lost |
| Putative cysteine-rich receptor-like protein kinase 30 | AT4G11460 | WT | frameshift variant |
| RING-H2 finger protein ATL17 | AT4G15975 | WT | frameshift variant |
| Disease resistance protein (TIR-NBS-LRR class) family | AT4G19520 | WT | frameshift variant |
| Toll-Interleukin-Resistance (TIR) domain family protein | AT4G19920 | WT | frameshift variant |
| Cyclic nucleotide-gated ion channel 2 | AT5G15410 | WT | frameshift variant |
| Probable glucan endo-1,3-beta-glucosidase BG5 | AT5G20340 | WT | frameshift variant |
| Disease resistance protein (TIR-NBS-LRR class) family | AT5G41740 | WT | frameshift variant |
| Disease resistance protein RPP8 | AT5G43470 | WT | frameshift variant |

**Supplementary Table 1.** Defense-annotated genes differentially mutated to high effect in the combined Landsberg accessions (La-0, L*er*-0, L*er*-1) vs Di-G. Defense annotation obtained from GO terms for Biological Processes.

| | | | | | | site class: 0 | | | site class: 1 | | | site class: 2a | | | site class: 2b | | | | |
|---|---|---|---|---|---|---|---|---|---|---|---|---|---|---|---|---|---|---|---|
| name | Model | Site-class model | fixed ω? | tree | ω | proportion | background ω | foreground ω | proportion | background ω | foreground ω | proportion | background ω | foreground ω | proportion | background ω | foreground ω | lnL | np |
| CYP82Cm0 | One Dn/Ds ratio | One w | no | ((((At82C2,Al8 | 0.208 | n/a | n/a | n/a | n/a | n/a | n/a | n/a | n/a | n/a | n/a | n/a | n/a | -5484.6177 | 15 |
| CYP82Cm1a | One Dn/Ds ratio | Neutral | no | ((((At82C2,Al8 | n/a | 0.8087 | 0.11791 | 0.11791 | 0.1913 | 1.00000 | 1.00000 | n/a | n/a | n/a | n/a | n/a | n/a | -5465.052 | 16 |
| CYP82C2C3_fixed | Several Dn/Ds ratios for branches | Selection | ω=1 | ((((At82C2,Al8 | n/a | 0.58993 | 0.11193 | 0.11193 | 0.13639 | 1.00000 | 1.00000 | 0.22228 | 0.11193 | 1.00000 | 0.05139 | 1.00000 | 1.00000 | -5462.967 | 17 |
| CYP82C2_fixed | Several Dn/Ds ratios for branches | Selection | ω=1 | ((((At82C2 #1, | n/a | 0.44962 | 0.11447 | 0.11447 | 0.08694 | 1.00000 | 1.00000 | 0.38835 | 0.11447 | 1.00000 | 0.07509 | 1.00000 | 1.00000 | -5461.548 | 17 |
| CYP82C2C3_unfixed | Several Dn/Ds ratios for branches | Selection | no | ((((At82C2,Al8 | n/a | 0.79502 | 0.11229 | 0.11229 | 0.18474 | 1.00000 | 1.00000 | 0.01643 | 0.11229 | 15.98222 | 0.00382 | 1.00000 | 15.98222 | -5461.549 | 18 |
| CYP82C2_unfixed | Several Dn/Ds ratios for branches | Selection | no | ((((At82C2 #1, | n/a | 0.49280 | 0.11428 | 0.11428 | 0.09571 | 1.00000 | 1.00000 | 0.34457 | 0.11428 | 1.12877 | 0.06692 | 1.00000 | 1.12877 | -5461.540 | 18 |

| | | | | Bayes Empirical Bayes analysis: positively selected sites |
|---|---|---|---|---|
| | null model | 2ΔlnL | *P* | H 250 |
| CYP82C2C3_unfixed | m1a | 7.01 | 0.030 | 0.962* |
| | fixed ω | 2.84 | 0.092 | |
| CYP82C2_unfixed | m1a | 7.02 | 0.030 | |
| | fixed ω | 0.02 | 0.897 | |

**Supplementary Table 2.** PAML output for purifying and positive selection on CYP82C family.

| **name** | **IDs** | **TE ID** | **identity to *EPCOT3*** |
|---|---|---|---|
| *RIX_Atal_Ta22_EPCOT3* | Chr4:15460121..15460384 | - | - |
| *RIX_Atal_Ta22_EPL1* | *AT4G29090* | *AT4TE67780* | 85.40% |
| *RIX_Atal_Ta22_EPL2* | *AT2G34320* | - | 67.00% |
| *RIX_Atal_Ta22_Ta22L1* | *AT2G45230* | *AT2TE84695* | 61.80% |
| *RIX_Atal_Ta22_Ta22L2* | *AT5G38285* | *AT5TE55355* | 61.10% |
| *RIX_Atal_Ta22_Ta22L3* | *AT5G42965* | - | 61.10% |
| *RIX_Atal_Ta22_Ta22L4* | Chr1:20156679..20156902 | - | 61.60% |
| *RIX_Atal_Ta22_Ta22* | AT5G28523 | - | 57.70% |
| *RIX_Atal_Ta22_Ta22L5* | Chr2:11595059..11595299 | - | 56.30% |
| *RIX_Atal_Ta22_Ta22L6* | Chr4:2716500..2717315 | *AT4TE12660* | 55.30% |
| *RIX_Atal_Ta22_Ta22L7* | *AT5G38920* | - | 61.50% |
| *RIX_Atal_Ta22_Ta22L8* | *AT2G16080* | - | 62.30% |

**Supplementary Table 3.** Characteristics of *EPCOT3*, *Ta22*, and related TE fragments identified in this study.

| | Di-G | Di-G | Di-G | Di-G | Di-G | Kz-13 | |
|---|---|---|---|---|---|---|---|
| SNP type | Ler-1 | Ler-0 | La-0 | Kz-13 | Zdrl-29 | Zdrl-29 | notes |
| A>C | 5.30 | 5.34 | 5.24 | 5.43 | 5.45 | 5.44 | |
| A>G | 13.46 | 13.58 | 12.71 | 12.83 | 13.43 | 13.16 | (transition) |
| A>T | 9.59 | 9.07 | 8.37 | 8.62 | 8.87 | 8.58 | |
| C>A | 5.45 | 5.41 | 5.34 | 5.49 | 5.46 | 5.41 | |
| C>G | 3.18 | 3.33 | 3.53 | 3.56 | 3.58 | 3.60 | |
| C>T | 13.30 | 13.56 | 14.91 | 14.15 | 13.36 | 13.92 | (transition) |
| G>A | 13.26 | 13.29 | 14.86 | 14.01 | 13.26 | 13.75 | (transition) |
| G>C | 3.18 | 3.38 | 3.62 | 3.63 | 3.48 | 3.64 | |
| G>T | 5.38 | 5.39 | 5.40 | 5.48 | 5.43 | 5.46 | |
| T>A | 9.67 | 9.14 | 8.31 | 8.70 | 8.97 | 8.57 | |
| T>C | 12.96 | 13.23 | 12.52 | 12.77 | 13.31 | 13.08 | (transition) |
| T>G | 5.27 | 5.29 | 5.19 | 5.32 | 5.39 | 5.40 | |
| **% transitions** | **52.98** | **53.65** | **55.00** | **53.76** | **53.37** | **53.91** | |
| indel size (nt) | | | | | | | |
| 2 | 64.14 | 67.74 | 67.74 | 66.62 | 68.66 | 70.50 | |
| 3 | 15.57 | 15.18 | 15.18 | 15.06 | 15.10 | 14.61 | |
| 4 | 7.43 | 6.74 | 6.74 | 7.15 | 6.80 | 6.39 | |
| 5 | 4.80 | 4.04 | 4.04 | 4.23 | 3.87 | 3.55 | |
| 6 | 2.33 | 1.78 | 1.78 | 2.04 | 1.66 | 1.71 | |
| 7 | 1.69 | 1.28 | 1.28 | 1.44 | 1.12 | 1.04 | |
| 8 | 1.19 | 0.95 | 0.95 | 1.02 | 0.78 | 0.71 | |
| 9 | 1.03 | 0.83 | 0.83 | 0.90 | 0.77 | 0.58 | |
| 10 | 0.57 | 0.52 | 0.52 | 0.53 | 0.47 | 0.30 | |
| 11 | 0.41 | 0.31 | 0.31 | 0.33 | 0.27 | 0.19 | |
| 12 | 0.31 | 0.22 | 0.22 | 0.24 | 0.17 | 0.16 | |
| 13 | 0.20 | 0.18 | 0.18 | 0.17 | 0.14 | 0.08 | |
| 14 | 0.11 | 0.09 | 0.09 | 0.09 | 0.07 | 0.05 | |
| 15 | 0.10 | 0.07 | 0.07 | 0.09 | 0.08 | 0.05 | |
| 16 | 0.033 | 0.020 | 0.020 | 0.031 | 0.018 | 0.023 | |
| 17 | 0.029 | 0.017 | 0.017 | 0.022 | 0.012 | 0.015 | |
| 18 | 0.017 | 0.010 | 0.010 | 0.016 | 0.009 | 0.010 | |
| 19 | 0.006 | 0.005 | 0.005 | 0.008 | 0.006 | 0.004 | |
| 20 | 0.015 | 0.000 | 0.000 | 0.008 | 0.000 | 0.008 | |
| 21 | 0.004 | 0.000 | 0.000 | 0.006 | 0.000 | 0.006 | |
| 22 | 0.002 | 0.000 | 0.000 | 0.000 | 0.000 | 0.000 | |
| 23 | 0.008 | 0.002 | 0.002 | 0.004 | 0.003 | 0.002 | |
| 24 | 0.004 | 0.000 | 0.000 | 0.002 | 0.000 | 0.002 | |
| 29 | 0.000 | 0.000 | 0.000 | 0.002 | 0.000 | 0.002 | |
| **% indels 10-29 nt** | **1.82** | **1.44** | **1.44** | **1.53** | **1.24** | **0.91** | |
| **% indels 15-29 nt** | **0.22** | **0.13** | **0.13** | **0.19** | **0.13** | **0.12** | |

**Supplementary Table 4.** Percentages of unique polymorphisms between two given ecotype pairs.

| Species | Accession/Cultivar | Mutant name | T-DNA/Stock Identifier | Source |
|---|---|---|---|---|
| *Arabidopsis lyrata* | | | CS22696 | Arabidopsis Biological Resource Center |
| *Arabidopsis thaliana* | Col-0 | *cyp82C2-2* | GABI_261D12 | Arabidopsis Biological Resource Center |
| *Arabidopsis thaliana* | Col-0 | *fls2-c* | SAIL_691_C4 | Arabidopsis Biological Resource Center |
| *Arabidopsis thaliana* | Col-0 | *rpm1-3* | CS8637 | Arabidopsis Biological Resource Center |
| *Arabidopsis thaliana* | Col-0 | *wrky33-1* | SALK_006603 | Jakub Rajniak; Arabidopsis Biological Resource Center |
| *Arabidopsis thaliana* | Col-0 | *cyp79B2 cyp79B3* | | JL Celenza |
| *Arabidopsis thaliana* | Be-0 | | CS964 | Arabidopsis Biological Resource Center |
| *Arabidopsis thaliana* | Chi-0 | | CS1072 | Arabidopsis Biological Resource Center |
| *Arabidopsis thaliana* | Chi-1 | | CS1074 | Arabidopsis Biological Resource Center |
| *Arabidopsis thaliana* | Col-0 | | CS1092 | Arabidopsis Biological Resource Center |
| *Arabidopsis thaliana* | Ct-1 | | CS22639 | Arabidopsis Biological Resource Center |
| *Arabidopsis thaliana* | Di-G | | CS910 | Arabidopsis Biological Resource Center |
| *Arabidopsis thaliana* | En-1 | | CS1136 | Arabidopsis Biological Resource Center |
| *Arabidopsis thaliana* | En-D | | CS920 | Arabidopsis Biological Resource Center |
| *Arabidopsis thaliana* | Est-1 | | CS39287 | Arabidopsis Biological Resource Center |
| *Arabidopsis thaliana* | Gr-1 | | CS1198 | Arabidopsis Biological Resource Center |
| *Arabidopsis thaliana* | Halca-1 | | CS76909 | Arabidopsis Biological Resource Center |
| *Arabidopsis thaliana* | Her-12 | | CS76920 | Arabidopsis Biological Resource Center |
| *Arabidopsis thaliana* | Kas-1 | | CS28376 | Arabidopsis Biological Resource Center |
| *Arabidopsis thaliana* | Kil-0 | | CS1270 | Arabidopsis Biological Resource Center |
| *Arabidopsis thaliana* | Ler-0 | | CS20 | Arabidopsis Biological Resource Center |
| *Arabidopsis thaliana* | Ler-1 | | CS22686 | Arabidopsis Biological Resource Center |
| *Arabidopsis thaliana* | Ma-0 | | CS1356 | Arabidopsis Biological Resource Center |
| *Arabidopsis thaliana* | Mc-0 | | CS1362 | Arabidopsis Biological Resource Center |
| *Arabidopsis thaliana* | Mir-0 | | CS1378 | Arabidopsis Biological Resource Center |
| *Arabidopsis thaliana* | Nc-1 | | CS1388 | Arabidopsis Biological Resource Center |
| *Arabidopsis thaliana* | Nd-0 | | CS1930 | Arabidopsis Biological Resource Center |
| *Arabidopsis thaliana* | No-0 | | CS24239 | Arabidopsis Biological Resource Center |
| *Arabidopsis thaliana* | Ob-0 | | CS38905 | Arabidopsis Biological Resource Center |
| *Arabidopsis thaliana* | Oy-0 | | CS22658 | Arabidopsis Biological Resource Center |
| *Arabidopsis thaliana* | Ped-0 | | CS76415 | Arabidopsis Biological Resource Center |
| *Arabidopsis thaliana* | Rel-0 | | CS77290 | Arabidopsis Biological Resource Center |
| *Arabidopsis thaliana* | RLD-1 | | CS913 | Arabidopsis Biological Resource Center |
| *Arabidopsis thaliana* | Sei-0 | | CS1504 | Arabidopsis Biological Resource Center |
| *Arabidopsis thaliana* | Sg-1 | | CS1518 | Arabidopsis Biological Resource Center |
| *Arabidopsis thaliana* | TDr-1 | | CS77345 | Arabidopsis Biological Resource Center |
| *Arabidopsis thaliana* | Tul-0 | | CS1570 | Arabidopsis Biological Resource Center |
| *Arabidopsis thaliana* | Ty-0 | | CS1572 | Arabidopsis Biological Resource Center |
| *Arabidopsis thaliana* | Ws-2 | | CS2360 | Arabidopsis Biological Resource Center |
| *Boechera holboelii* | 910 | | | T Mitchell-Olds |
| *Brassica rapa* | Mizuna | | | Stover Seeds (Sun Valley, CA) |
| *Capsella rubella* | Monte Gargano | | CS22697 | Arabidopsis Biological Resource Center |
| *Cardamine hirsuta* | | | | field site at 41˚46'24"N 70˚03'03"W |
| *Crucihimalaya lasiocarpa* | | | CS6191 | Arabidopsis Biological Resource Center |
| *Erysimum chieri* | | | | J.L. Hudson, Seedsman (La Honda, CA) |
| *Olimarabidopsis cabulica* | | | CS4653 | Arabidopsis Biological Resource Center |
| *Polyctenium fremontii* | | | | Kew Gardens (London, UK) |

**Supplementary Table 5.** Seed stocks used in this study.

| | TAIR ID | gene | primers | sequence | efficiency | R2 for efficiency |
|---|---|---|---|---|---|---|
| qPCR | AT2G30750 | CYP71A12 | 71A12R | GATTATCACCT CGGTTCCT | 93.1 | 0.998 |
| qPCR | AT2G30750 | CYP71A12 | 71A12F | CCACTAATACTT CCCAGATTA | 93.1 | 0.998 |
| qPCR | AL4G25660 | AlCYP71A12 | Al71A12_L1 | CCACTAATACTT CCTAGACTA | 100 | |
| qPCR | AL4G25660 | AlCYP71A12 | Al71A12_R1 | GATTATAACCTC TGTCCCT | 100 | |
| qPCR | AT2G30770 | CYP71A13 | 71A13F | GCTTCGGTTGC ATCCTTCT | 87.5 | 0.996 |
| qPCR | AT2G30770 | CYP71A13 | 71A13R | ATTGATTATCAC CTCTGTCC | 87.5 | 0.996 |
| qPCR | AT3G26830 | CYP71B15/PAD3 | PAD3-L1 | ACGAGCATCTT AAGCCTGGA | 78.4 | 0.998 |
| qPCR | AT3G26830 | CYP71B15/PAD3 | PAD3-R1 | TCGGTCATTCC CCATAGTGT | 78.4 | 0.998 |
| qPCR | AT4G39950 | CYP79B2 | 79B2-1R | GTAACTTCGGA GCATTCGT | 95 | 1 |
| qPCR | AT4G39950 | CYP79B2 | 79B2-2F | TCGCCGGATAT CACATCC | 95 | 1 |
| qPCR | AL7G12200 | AlCYP79B2 | Al79B2_R1 | GTGACTTCGGA GCATTCGT | 100 | |
| qPCR | AT2G22330 | CYP79B3 | 79B3-1R | AGTCACTTCCG AACACTCA | 92.8 | 0.999 |
| qPCR | AT2G22330 | CYP79B3 | 79B3-2F | TCGCAGGTTAC CATATTCC | 92.8 | 0.999 |
| qPCR | AT5G57220 | CYP81F2 | 81F2-F | CTCATGCTCAG TATGATGC | 86.2 | 0.997 |
| qPCR | AT5G57220 | CYP81F2 | 81F2-2R | CTCCAATCTTCT CGTCTATC | 86.2 | 0.997 |
| qPCR | AT4G31970 | CYP82C2 | 82C2-L1 | CAAGCATGTCC GTGTTTCTG | 91.6 | 1 |
| qPCR | AT4G31970 | CYP82C2 | 82C2-R1 | GCATCTTCAGG GGATAACGA | 91.6 | 1 |
| qPCR | AL7G19940 | AlCYP82C2 | Al82C2_L1 | TGAGATCGCAA TGGGTGTG | 100 | |
| qPCR | AL7G19940 | AlCYP82C2 | Al82C2_R1 | TCTTTATCCTCG GAAGATAC | 100 | |
| qPCR | AT3G31940 | CYP82C4 | At82C4_L1 | TGAGATCACAA TGGGTGTG | 100 | 0.998 |
| qPCR | AT3G31940 | CYP82C4 | At82C4_R1 | TCCGACGATAC TGAGCC | 100 | 0.998 |
| qPCR | AL7G19950 | AlCYP82C4 | Al82C4_L1 | TGAGATCTCAA TGGGTGTG | 100 | |
| qPCR | AL7G19950 | AlCYP82C4 | Al82C4_R1 | CTCCGAGGATT GTGAGC | 100 | |
| qPCR | AT4G31500 | CYP83B1 | SUR2-1R | TCACGCCATAT CTACCAGC | 87.3 | 1 |
| qPCR | AT4G31500 | CYP83B1 | SUR2-2F | TGGACGTCATG ACTGGAC | 87.3 | 1 |

| | | | | | | |
|---|---|---|---|---|---|---|
| qPCR | AT3G13920 | eIF4A1 | EIF4AF | TCTGCACCAGAAGGCACA | 100 | |
| qPCR | AT3G13920 | eIF4A1 | EIF4A2R | TCATAGGATGTGAAGAACTC | 100 | |
| qPCR | AL3G26100 | AlEI4A1 | AlEI4A1-R | TCATAAGATGTGAAGAACTC | 100 | |
| qPCR | AT3G13610 | F6'H1 | F6'H-2F | CTGATATCTCAGGAATGAAACG | 100 | 0.997 |
| qPCR | AT3G13610 | F6'H1 | F6'H-2R | GGTAGTAGTTAAGGTTGACTC | 100 | 0.997 |
| qPCR | AT1G26380 | FOX1 | FOX1-1F | ATGATGGATCGGATTCCGT | 98.5 | 0.999 |
| qPCR | AT1G26380 | FOX1 | FOX1-2R | ACTCAGGTTACTCTCTGTG | 98.5 | 0.999 |
| qPCR | AL1G39380 | AlFOX1 | AlFOX1_L1 | ATGATGGACCGGATTCCGT | 100 | |
| qPCR | AT2G20610 | SUR1 | SUR1-F | TGTAACACCTTGGTGGTCT | 91.5 | 1 |
| qPCR | AT2G20610 | SUR1 | SUR1-R | CCTGTATTGACACTAGCAG | 91.5 | 1 |
| qPCR | AT2G38470 | WRKY33 | WRKY33-1F | GAAGCAAAGAGATGGAAAGG | 99.4 | 1 |
| qPCR | AT2G38470 | WRKY33 | WRKY33-1R | CTACGATTCTCGGCTCTC | 99.4 | 1 |
| | TAIR ID | gene | primers | sequence | amplicon | |
| ChIP-PCR | AT3G26830 | CYP71B15 | 71B15pr4F | gaggtgtgcaatatggac | W3 | |
| ChIP-PCR | AT3G26830 | CYP71B15 | 71B15pr4R | attaggttgcgctgacca | W3 | |
| ChIP-PCR | AT3G26830 | CYP71B15 | 71B15pr3F | ataccctatcatgttttactc | W2 | |
| ChIP-PCR | AT3G26830 | CYP71B15 | 71B15pr3R | agtcaacaattttagcggtg | W2 | |
| ChIP-PCR | AT3G26830 | CYP71B15 | 71B15pr2F | gaagaagttttggaatagcc | W1 | |
| ChIP-PCR | AT3G26830 | CYP71B15 | 71B15pr2R | agtcaatctttttgtatcatgc | W1 | |
| ChIP-PCR | AT1G26380 | FOX1 | FOX1pr1F | gtattttacaacaacgtagcc | W2 | |
| ChIP-PCR | AT1G26380 | FOX1 | FOX1pr1R | tatctactaattgaaataattgcg | W2 | |
| ChIP-PCR | AT1G26380 | FOX1 | FOX1pr2F | gtactatacgtattacgtacg | W1 | |
| ChIP-PCR | AT1G26380 | FOX1 | FOX1pr2R | ctttcattttgctgtttgg | W1 | |
| ChIP-PCR | AT4G31970 | CYP82C2 | 82C2pr1F | acagaaaaacccactaaagtc | W5 | |
| ChIP-PCR | AT4G31970 | CYP82C2 | 82C2pr1R | ggtttgtattgattttgttagtc | W5 | |
| ChIP-PCR | AT4G31970 | CYP82C2 | 82C2pr2F | cttgcccaactaggcac | W4 | |

| | | | | | |
|---|---|---|---|---|---|
| ChIP-PCR | AT4G31970 | CYP82C2 | 82C2pr2R | atgatctgcagaggttgct | W4 |
| ChIP-PCR | AT4G31970 | CYP82C2 | 82C2pr3F | cccataactcagagctcc | W3 |
| ChIP-PCR | AT4G31970 | CYP82C2 | 82C2pr3R | aattatattttttgtttttgtatgag | W3 |
| ChIP-PCR | AT4G31970 | CYP82C2 | 82C2pr7F | atttgagttattattatgaattcag | W2 |
| ChIP-PCR | AT4G31970 | CYP82C2 | 82C2pr7R | tttccaccatttatcttaacca | W2 |
| ChIP-PCR | AT4G31970 | CYP82C2 | 82C2pr5F | ttttgaccatatacaatttgcg | W1 |
| ChIP-PCR | AT4G31970 | CYP82C2 | 82C2pr5R | tatacacacacatattatcggt | W1 |
| ChIP-PCR | AT4G29090 | EPL1 | 4G29090pr1F | ctagcccaactaggcac | W |
| ChIP-PCR | AT4G29090 | EPL1 | 4G29090pr1R | aagatctgcagaggttgct | W |
| ChIP-PCR | AT2G34320 | EPL2 | 2G34320pr2F | CCTTCAACCGGCTTTGG | W |
| ChIP-PCR | AT2G34320 | EPL2 | 2G34320pr2R | AATCAAAGTGCTTCTAAGCC | W |
| | TAIR ID | gene | primers | sequence | notes |
| RT-PCR | AT2G38470 | WRKY33 | AtWRKY33-1F | GAAGCAAAGAGATGGAAAGG | |
| RT-PCR | AT2G38470 | WRKY33 | AtWRKY33-1R | CTACGATTCTCGGCTCTC | |
| RT-PCR | AT4G31970 | CYP82C2 | AtCYP82C2-L2 | TCTGAGATCTCAATGGTTATGC | (downstream of cyp82C2-2 T-DNA insertion) |
| RT-PCR | AT4G31970 | CYP82C2 | AtCYP82C2-R3 | GCATCTTCAGGGGATAACGA | (downstream of cyp82C2-2 T-DNA insertion; used to amplify AtCYP82C2 and AtCYP82C2-long) |
| RT-PCR | AT4G31970 | CYP82C2 | AtCYP82C2-1F | TGCACCAAGTGGTGCGTG | (upstream of cyp82C2-2 T-DNA insertion; used only to amplify AtCYP82C2-long) |
| RT-PCR | AL7G19940 | AlCYP82C2 | AlCYP82C2-L1 | TGAGATCGCAATGGGTGTG | |
| RT-PCR | AL7G19940 | AlCYP82C2 | AlCYP82C2-R3 | CCGAGGATAAAGAGCCG | |
| RT-PCR | | NbActin1 | NbActin1-F | ATCCTCACAGAGCGTGGTTAC | |
| RT-PCR | | NbActin1 | NbActin1-R | CACTGAGCACTATGTTTCCGT | |

**Supplementary Table 6.** Primer sequences used in this study.

| Glucosinolates | | | |
|---|---|---|---|
| retention (min) | flow (mL/min) | %A (H2O) | %B (90% acetonitrile) |
| 0 | 0.5 | 98 | 2 |
| 1 | 0.5 | 98 | 2 |
| 6 | 0.5 | 94 | 6 |
| 8 | 0.5 | 92 | 8 |
| 16 | 0.5 | 77 | 23 |
| 20 | 0.5 | 69 | 31 |
| 28 | 0.5 | 0 | 100 |
| 33 | 0.5 | 0 | 100 |
| 34 | 0.5 | 98 | 2 |
| 44 | 0.5 | 98 | 2 |

| Camalexin/ICN/4OH-ICN | | | | |
|---|---|---|---|---|
| retention (min) | flow (mL/min) | %A (0.005% [v/v] formic acid) | %B (methanol + 0.005% [v/v] formic acid) | %C (90% acetonitrile + 0.005% [v/v] formic acid) |
| 0 | 0.5 | 65 | 15 | 20 |
| 5 | 0.5 | 60 | 20 | 20 |
| 10 | 0.5 | 54.3 | 25.7 | 20 |
| 13 | 0.5 | 53.1 | 26.9 | 20 |
| 13.5 | 0.5 | 52.5 | 27.5 | 20 |
| 16 | 0.5 | 52.5 | 27.5 | 20 |
| 16 | 0.5 | 0 | 80 | 20 |
| 21 | 0.5 | 0 | 80 | 20 |
| 21.5 | 0.5 | 65 | 15 | 20 |
| 26.5 | 0.5 | 65 | 15 | 20 |

**Supplementary Table 7.** LC gradients used in this study.

| species | genome release | chromsome/contig(s) for CYP82C analysis | source |
|---|---|---|---|
| *Arabidopsis thaliana* | v9 | chr4 | Phytozome 12 (Goodstein et al., 2011) |
| *Arabidopsis halleri* | v1.1 | Scaffold3172 | Phytozome 12 (Goodstein et al., 2011) |
| *Arabidopsis lyrata* | v2.1 | scaffold_7 | Phytozome 12 (Goodstein et al., 2011) |
| *Capsella rubella* | v1.0 | scaffold_7 | Phytozome 12 (Goodstein et al., 2011) |
| *Capsella grandiflora* | v1.1 | Scaffold3807, Scaffold12385 | Phytozome 12 (Goodstein et al., 2011) |
| *Boechera stricta* | v1.2 | Scaffold7867 | Phytozome 12 (Goodstein et al., 2011) |
| Notes: The coding sequence for CgCYP82C2 spans two scaffold sequences. This region was manually translated and assembled, resulting in a gap of a single amino acid, which was annotated as “X”. AhCYP82C2 was excluded from protein sequence analyses due to a 387nt gap in the third exon. The 2,072nt gap upstream of *CYP82C4* was removed for mVISTA alignments. | | | |

**Supplementary Table 8.** Contigs and chromosomes used for CYP82C analysis.
